# Activation and *in vivo* evolution of the MAIT cell transcriptome in mice and humans reveals diverse functionality

**DOI:** 10.1101/490649

**Authors:** Timothy SC Hinks, Emanuele Marchi, Maisha Jabeen, Moshe Olshansky, Ayako Kurioka, Troi J Pediongco, Bronwyn S Meehan, Lyudmila Kostenko, Stephen J Turner, Alexandra J Corbett, Zhenjun Chen, Paul Klenerman, James McCluskey

**Affiliations:** Department of Microbiology and Immunology, Peter Doherty Institute for Infection and Immunity, University of Melbourne, Parkville, Victoria 3000, Australia.; Respiratory Medicine Unit and National Institute for Health Research (NIHR) Oxford Biomedical Research Centre (BRC), Nuffield Department of Medicine Experimental Medicine, University of Oxford, OX3 9DU, Oxfordshire, UK; Peter Medawar Building for Pathogen Research and Translational Gastroenterology Unit, Nuffield Department of Clinical Medicine, University of Oxford, OX3 3SY; Infection and Immunity Program and The Department of Biochemistry and Molecular Biology, Biomedicine Discovery Institute, Monash University, Clayton, Australia.

**Author notes:** Corresponding author and lead contact: Dr TSC Hinks, Respiratory Medicine Unit, NDM Experimental Medicine, University of Oxford, Level 7, John Radcliffe Hospital, Oxford, OX3 9DU; @HinksLab.

**Keywords:** Mucosal-associated invariant T cell, T cell, transcriptome, MHC-related protein 1, activation, lung, human, mouse, riboflavin

## Abstract

Mucosal-associated invariant T (MAIT) cells are MR1-restricted innate-like T cells conserved across mammalian species, including mice and humans. By sequencing RNA from sorted MR1-5-OP-RU tetramer^+^ cells derived from either human blood or murine lungs, we define the basic transcriptome of an activated MAIT cell in both species and demonstrate how this profile changes during resolution and reinfection phases of infection. We observe strong similarities between MAIT cells in humans and mice. Compared with previously published T cell transcriptomes, MAIT cells displayed most similarity to iNKT cells when activated, but to γδ T cells, after resolution of infection. In both species activation leads to strong expression of pro-inflammatory cytokines and chemokines, and also a strong tissue repair signature, recently described in murine commensal-specific H2-M3-restricted T cells. These data define the requirements for, and consequences of, MAIT cell activation, revealing a tissue repair phenotype expressed upon MAIT cell activation in both species.

---

Mucosal-associated invariant T (MAIT) cells are innate-like T cells which express a ‘semi-invariant’ αβ T cell receptor (TCR) and recognise metabolic derivatives of riboflavin biosynthesis^**1–3**^ presented on the restriction molecule major histocompatibility complex (MHC)-related protein-1 (MR1)^**4,5**^. These antigens, which include the potent MAIT cell ligand 5-(2-oxopropylideneamino)-6-D-ribitylaminouracil (5-OP-RU)^**6**^, are produced by a wide variety of bacteria, mycobacteria and yeasts^**1,7**^, but are absent from mammals, and therefore allows host – pathogen discrimination. MAIT cells have a strong pro-inflammatory phenotype, and produce interferon-γ (IFN-γ), TNF and IL-17A after phorbol myristate acetate (PMA) and ionomycin stimulation^**8**^.

Whilst baseline frequencies of MAIT cells are low in specific-pathogen free C57BL/6 mice, we, and others, have previously shown that MAIT cells can be activated and expand *in vivo* in response to pulmonary infection with specific intracellular bacteria expressing the riboflavin pathway – *Salmonella* Typhimurium^9^, *Legionella spp*^*10*^, and *Francisella tularensis*^11,12^ – or in response to synthetic 5-OP-RU accompanied by a Tolllike receptor agonst^9^, providing valuable models to dissect MAIT cell biology.

To date the requirements for TCR-dependent activation of MAIT cells *in vivo* have not been systematically characterised, nor have the consequences of such activation been fully defined. Here we have used MR1 tetramers^2^ loaded with 5-OP-RU to specifically identify MAIT cells from human peripheral blood and murine lungs, allowing us to assess the requirements for, and consequences of, MAIT cell activation *ex vivo* and *in vivo*. Using a transcriptomic approach on sorted MR1-5-OP-RU tetramer^+^ cells we define the transcriptome of an activated MAIT cell in both species and explore how this changes during the resolution and reinfection phases of infection.

Our data reveal strong similarities between MAIT cells in humans and in mice at a transcriptional level, show that MAIT cells displayed the closest similarities to invariant natural killer T (iNKT) cells when activated, but after resolution of infection were more comparable to γδ T cells, and reveal a previously unknown tissue repair phenotype expressed upon MAIT cell activation in both species.

## Results

### ctivation requirements of MAIT cells *in vivo*

First we aimed to test, systematically, the activation requirements of MAIT cells *in vivo* in mouse lungs. We have previously shown that pulmonary MAIT cell frequencies in mice can be markedly enhanced by intranasal administration of 5-OP-RU if it is co-administered with S-[2,3-bis(palmitoyloxy)propyl] cysteine (Pam2Cys), CpG ODN 1668 or polyinosinic:polycytidylic acid (poly I:C), which are agonists for TLR2/6, TLR9 and TLR3, respectively. We therefore investigated agonists for each of the murine TLRs, using the maximum doses presented in a literature review of previous studies of these compounds. All animals received the relevant TLR intranasally on day 0. In experimental animals this was administered in combination with 76 pmol 5-OP-RU on day 0, with repeated inoculae of 76 pmol 5-OP-RU on days 1, 2 and 4. Control mice received the same TLR ligand and 76 pmol of the non-activating MR1 ligand 6-formyl pterin (6-FP) according to the same schedule, or the TLR ligand alone (Supplementary Table S1). We observed 15-180-fold enrichment of pulmonary CD3^+^CD45.2^+^CD19^-^MR1-5-OP-RU tetramer^+^ MAIT cell frequencies at day 7, after administration of 5-OP-RU with agonists of TLR3 (high molecular weight poly I:C), TLR4 (Lipopolysaccharide from *E.coli*), TLR2/6 (FSL-1 (Pam2CGDPKHPKSF)) and TLR9 (CpG ODN1826), but not with agonists of TLR1/2 (Pam3CSK4), TLR2 (heat killed *Listeria monocytogenes*), TLR5 (Flagellin from *S.typhimurium*), TLR7 (Imiquimod)(Figure 1), suggesting there is a specific and restricted range of danger signals that are capable of providing the necessary co-stimulus to drive MAIT cell accumulation in response to 5-OP-RU antigen.

**Figure 1.**
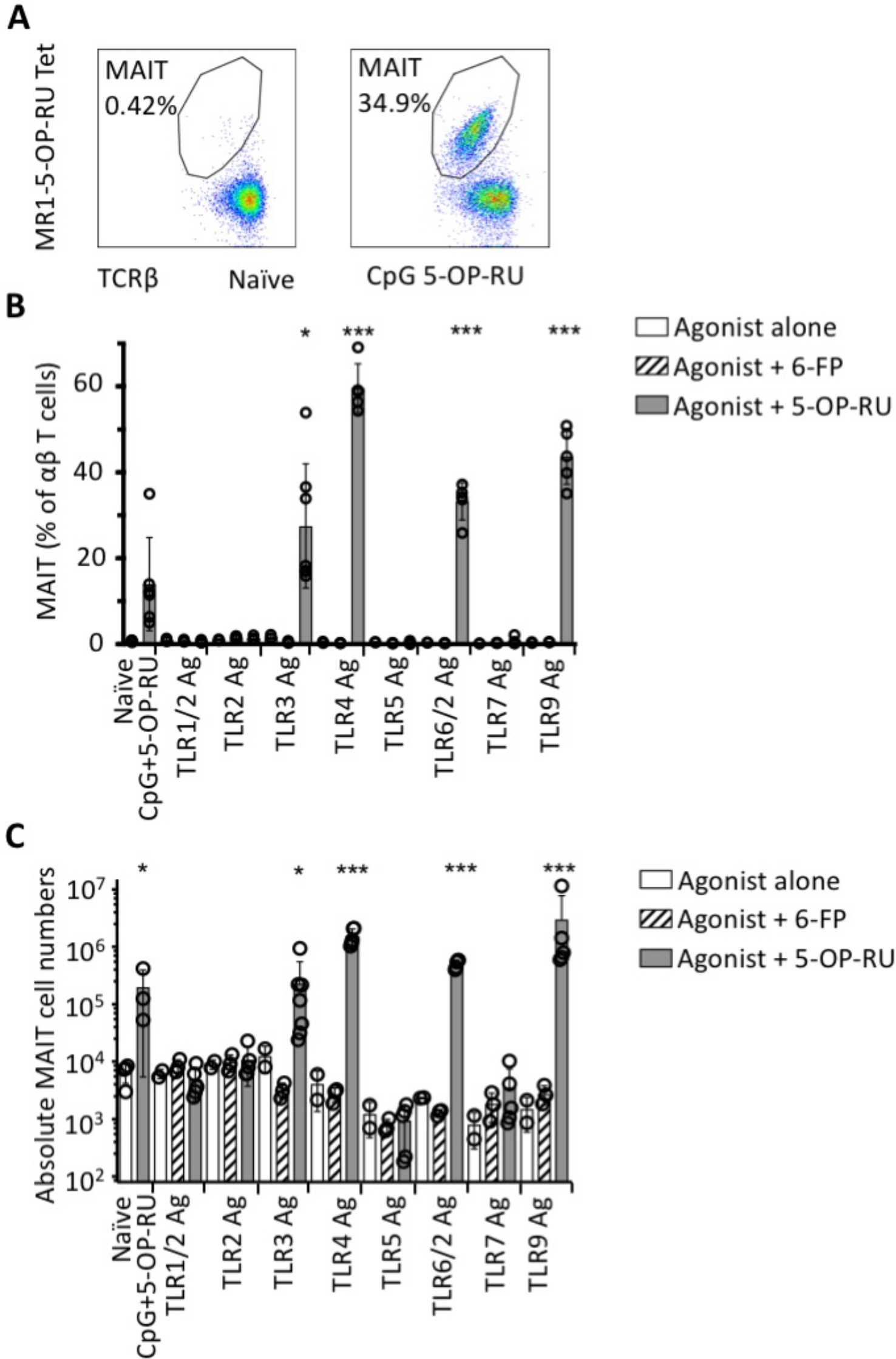
Costimulatory requirements for MAIT cell activation *in vivo*. **(A)** Representative flow-cytometry plots showing MAIT cell percentage among TCRβ^+^ lymphocytes in the lungs of C57BL/6 mice with or without prior stimulation with intranasal CpG and 5-OP-RU. Relative **(B)** and absolute **(C)** numbers of MR1-5-OP-RU tetramer^+^ MAIT cells in the lungs of C57BL/6 mice 7 days after intranasal exposure to specific TLR agonists either alone, or in combination with 76 pmol 6-FP, or with 76 pmol 5-OP-RU. Control mice received nothing (n=4, naïve) or CpG with 5-OP-RU (n=3). Experiments used n=5 (5-OP-RU treated), n=3 (6-FP treated) or n=2 (TLR agonist alone) mice per group. The experiment was subsequently repeated with similar results. Statistical tests: unpaired t tests, comparing TLR + 5-OP-RU with naïve control (n=4), on untransformed (B) or log-transformed (C) data, with Bonferroni corrections * P<0.05, *** P<0.001.

### Transcriptomic profile of activated human and murine MAIT cells

Having explored the requirements for activation of MAIT cells, we sought to describe in detail the consequences of their activation using a transcriptomic approach to define the basic transcriptome of a MAIT cell in both humans and mice and to determine how this is modulated by activation. Fresh human peripheral blood cells were obtained from three donors. These were cultured for 6 hours with (‘stimulated’) or without (‘unstimulated’) 10 nM 5-OP-RU, magnetically enriched on MR1-tetramer^+^ cells, and flow-sorted for RNA sequencing of live CD3^+^TCR Vα7.2^+^ MR1-5-OP-RU tetramer+ MAIT cells, and of unstimulated naïve live CD8^+^CD45RA^+^ T cells as a comparator cell type (**Table 1**).

**Table 1.**
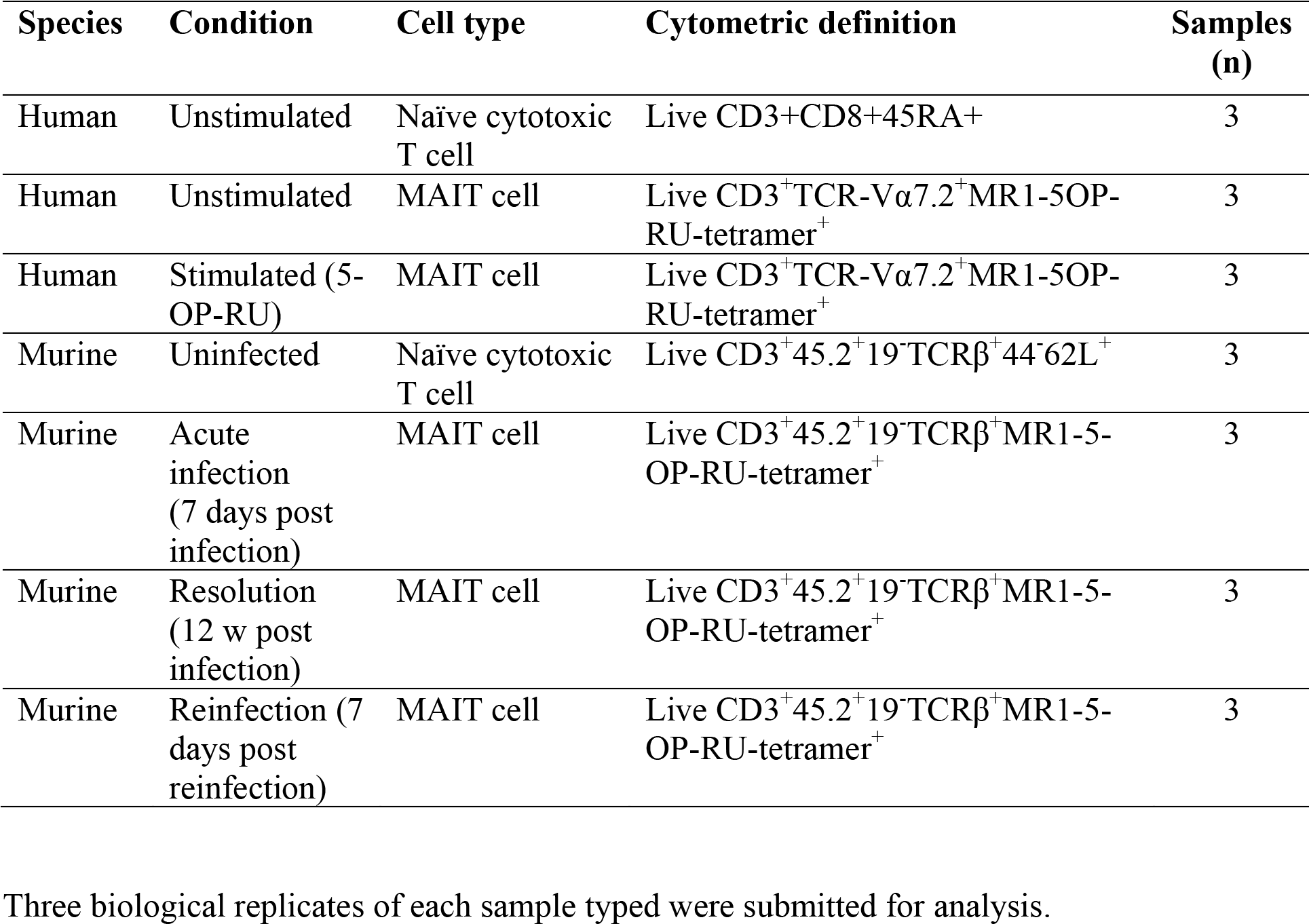
Definitions of transcriptomic samples.

We have previously shown that pulmonary infection of mice with the intracellular pathogen *Legionella longbeachae* induces strong TCR-mediated MAIT cell activation, and that this plays a significant role in host immune protection, thus constituting a physiologically relevant model of *in vivo* MAIT cell activation. Using this model, which induces a rapid and sustained expansion of MAIT cells in the lung (Supplementary figure S1) we therefore included within the same sequencing experiment live pulmonary CD3^+^45.2^+^19^-^MR1-5-OP-RU tetramer^+^ MAIT cells which were magnetically enriched and flow-sorted from the lungs of mice 7 days after infection with 1x10^4^ CFU *L. longbeachae* (‘acute’), or at least 12 weeks post infection (‘resolution’) or 7 days after a second intranasal infection with 2x10^4^ CFU *L. longbeachae* in mice that had recovered from infection 12 weeks previously (‘reinfection’). Live CD3^+^CD45.2^+^CD19^-^CD8^+^CD44^-^CD62L^+^ naïve T cells from uninfected mice were used as a comparator cell type.

The number of differentially expressed genes (DEG) in activated MAIT cells compared with naïve CD8^+^ T cells was 4613 genes in human 5-OP-RU-stimulated MAIT cells, and 3758 genes in acutely infected mice at a false discovery rate (FDR) p value of <0.05 and minimum log_(2)_ fold change of ±1 (Numbers of DEG are shown in Table 2; full lists of DEG are shown in Supplementary tables S2 and S3). These genes constitute the basic transcriptome of an activated MAIT cell in each species. To explore the nature of these gene profiles further we compared different activation states of MAIT cells. In humans 3227 genes were differentially expressed between stimulated and unstimulated MAIT cells and could therefore be considered the direct signature of TCR-mediated MAIT cell activation, whilst 968 genes were differentially expressed between unstimulated MAIT cells and naïve CD8^+^ T cells, and therefore are more related to constitutive differences between T cell lineages (Supplementary table S2). In mice 1889 genes were differentially expressed between acute infection and resolution of infection, analogous to the signature of TCR-mediated activation (Supplementary table S3).

**Table 2.** Numbers of differentially expressed genes

|  | Human |  |  | Murine |  |  |
| --- | --- | --- | --- | --- | --- | --- |
|  | Unstimulated<br>MAIT | Stimulated<br>MAIT | Stimulated<br>MAIT | Acute<br>infection<br>MAIT | Resolved<br>infection MAIT | Reinfection<br>MAIT |
|  | v<br>Unstimulated<br>CD8+45RA+ | v<br>Unstimulated<br>MAIT | v<br>Unstimulated<br>CD8+45RA+ | v<br>Uninfected<br>CD8<sup>+</sup>44<sup>Lo</sup>62<sup>Hi</sup> | v<br>Acute infection<br>MAIT | v<br>Acute infection<br>MAIT |
| Number of<br>upregulated genes | 411 | 1384 | 1651 | 1985 | 942 | 235 |
| Number of<br>downregulated genes | 557 | 1843 | 2512 | 1773 | 947 | 241 |

Genes shown are censored at FDR $P \leq 0.05$ and log(2) fold change of $\pm 1$ .

Analysis of TCR genes over-represented in MAIT cells confirmed highly significant selective use of TRAV1-2 with TRBV6-4, TRBV6-1 and TRBV20-1 in humans and Trbv13-5 with Trav1 and Traj33 in mice, in MAIT cells compared naïve CD8 T cells, as expected^4,5,13,14^ (Supplementary tables S4, S5).

Focussed analysis of known cytokine genes confirmed a strong upregulation of several pro-inflammatory type 1 and type 17 cytokines, especially CSF2 (GM-CSF), IL-17A, LIF, TNF, IFN-γ, and IL-17F) which were highly upregulated in both mouse and human in activated MAIT cells (Tables 3,4). Expression of selected cytokines was confirmed by flow cytometry (Supplementary figures S2, S3). Likewise there was significant, but more modest, upregulation of LTA (lymphotoxin A) and CSF1 (M-CSF) in both species. Some features were observed only in one species, notably IL-2 and TNSF14 (LIGHT) produced by activated human MAIT cells, and Tnfsf11 (TRANCE, RANKL) by murine MAIT cells. In contrast to activation-induced cytokines, expression of the anti-apoptotic cytokine IL-15, implicated in the development and maturation of memory CD8^+^ T cells^15^, was restricted to MAIT cells in their resting state: human unstimulated MAIT cells, or murine MAIT cells at resolution of infection. In mice resolution of infection was also associated specifically with strong expression of Tnfsf18 (GITRL), confirmed by flow cytometry. Expression of cytokine receptors is shown in Supplementary tables S6, S7 and of recognised ‘CD markers’ in Supplementary tables S8, S9.

**Table 3.** Differentially expressed cytokine genes. Human.

| Stimulated MAIT |  |  |  |  |  | Stimulated MAIT |  |  |  |  |  | Unstimulated MAIT |  |  |  |  |  |
| --- | --- | --- | --- | --- | --- | --- | --- | --- | --- | --- | --- | --- | --- | --- | --- | --- | --- |
| v |  |  |  |  |  | v |  |  |  |  |  | v |  |  |  |  |  |
| Unstimulated CD8+45RA+ |  |  |  |  |  | Unstimulated MAIT |  |  |  |  |  | Unstimulated CD8+45RA+ |  |  |  |  |  |
| Gene | Log fold change | Log CP M | LR | P value | FDR P value | Gene | Log fold change | Log CP M | LR | P value | FDR P value | Gene | Log fold change | Log CP M | LR | P value | FDR P value |
| CSF2 | 15 | 7.4 | 62 | 3E-15 | 0.00 | CSF2 | 15 | 7.4 | 64 | 1E-15 | 0.00 | LIF | 4.7 | 1.8 | 7.2 | 0.007 | 0.02 |
| IL17A | 12 | 4.9 | 74 | 8E-18 | 0.00 | IL17A | 12 | 4.9 | 77 | 1E-18 | 0.00 | TGFA | 4.4 | 3.3 | 43 | 6E-11 | 0.00 |
| IL2 | 11 | 4.0 | 42 | 1E-10 | 0.00 | IFNG | 9.7 | 8.4 | 61 | 6E-15 | 0.00 | TNF | 2.4 | 9.0 | 7.9 | 0.005 | 0.02 |
| LIF | 8.9 | 1.8 | 26 | 3E-07 | 0.00 | IL17F | 8.3 | 1.2 | 35 | 3E-09 | 0.00 | TNFS F13B | 2.3 | 3.3 | 9.5 | 0.002 | 0.01 |
| TNF | 8.7 | 9.0 | 59 | 2E-14 | 0.00 | IL2 | 8.0 | 4.0 | 33 | 1E-08 | 0.00 | TNFS F14 | 2.1 | 8.0 | 22 | 2E-06 | 0.00 |
| IFNG | 8.4 | 8.4 | 51 | 1E-12 | 0.00 | TNF | 6.3 | 9.0 | 38 | 8E-10 | 0.00 | CSF1 | 2.0 | 5.1 | 21 | 7E-06 | 0.00 |
| IL17F | 8.3 | 1.2 | 32 | 1E-08 | 0.00 | LIF | 4.2 | 1.8 | 12 | 0.0005 | 0.00 | IL15 | 1.9 | 2.3 | 5.8 | 0.02 | 0.04 |
| TNFSF14 | 5.3 | 8.0 | 113 | 2E-26 | 0.00 | IL10 | 4.1 | 1.5 | 14 | 0.0002 | 0.00 | IL24 | -1.8 | 2.4 | 6.2 | 0.01 | 0.04 |
| CSF1 | 5.2 | 5.1 | 120 | 7E-28 | 0.00 | TNFSF14 | 3.2 | 8.0 | 52 | 6E-13 | 0.00 | CXCR2 | -4.0 | 3.0 | 9.7 | 0.001 | 0.01 |
| IL26 | 4.6 | 1.8 | 16 | 7E-05 | 0.00 | CSF1 | 3.2 | 5.1 | 59 | 2E-14 | 0.00 | CXCR1 | -4.1 | 2.6 | 14 | 0.0001 | 0.00 |
| LTA | 3.3 | 7.0 | 48 | 4E-12 | 0.00 | LTA | 3.1 | 7.0 | 43 | 7E-11 | 0.00 |  |  |  |  |  |  |
| TGFA | 2.9 | 3.3 | 19 | 1E-05 | 0.00 | IL16 | -1.1 | 9.3 | 5.6 | 0.018 | 0.04 |  |  |  |  |  |  |
| TNFSF13B | 2.6 | 3.2 | 12 | 0.0004 | 0.00 | TGFA | -1.6 | 3.3 | 7.4 | 0.006 | 0.02 |  |  |  |  |  |  |

|  |  |  |  |  |  |  |  |  |  |  |  |
| --- | --- | --- | --- | --- | --- | --- | --- | --- | --- | --- | --- |
| IL23A | 2.3 | 4.7 | 12 | 0.0004 | 0.00 | IL32 | -1.6 | 9.7 | 20 | 8.1E-06 | 0.00 |
| IL32 | -1.2 | 9.7 | 12 | 0.0005 | 0.00 | IL18BP | -3.0 | 5.1 | 40 | 2.9E-10 | 0.00 |
| IL16 | -1.5 | 9.3 | 11 | 0.001 | 0.00 | CXCR1 | -5.7 | 2.6 | 13 | 0.0004 | 0.00 |
| IL24 | -2.0 | 2.4 | 7.7 | 0.005 | 0.01 |  |  |  |  |  |  |
| IL18BP | -3.6 | 5.1 | 57 | 6E-14 | 0.00 |  |  |  |  |  |  |
| CXCR2 | -4.9 | 3.0 | 13 | 0.0002 | 0.00 |  |  |  |  |  |  |
| CXCR1 | -9.8 | 2.6 | 37 | 1E-09 | 0.00 |  |  |  |  |  |  |

Genes shown are censored at FDR $P \leq 0.05$ and $\log(2)$ fold change of $\pm 1$ , and ordered by log fold change. Genes highlighted in bold are differentially expressed in both human and mouse. CPM, Counts per million; FDR, false discovery rate; LR, likelihood ratio.

**Table 4.** Differentially expressed cytokine genes. Mouse.

| Acute infection MAIT |  |  |  |  |  | Resolved infection MAIT cells |  |  |  |  |  |
| --- | --- | --- | --- | --- | --- | --- | --- | --- | --- | --- | --- |
| v<br>Uninfected CD8<sup>+</sup>44<sup>Lo</sup>62<sup>Hi</sup> |  |  |  |  |  | v<br>Acute infection MAIT cells |  |  |  |  |  |
| Gene | Log fold change | Log CPM | LR | P value | FDR P value | Gene | Log fold change | Log CPM | LR | P value | FDR P value |
| **Il17a** | 14 | 7.3 | 316 | 9E-71 | 1E-68 | Tnfsf18 (GITRL) | 5.9 | -0.52 | 14 | 0.0002 | 0.001 |
| **Ifng** | 12 | 5.9 | 222 | 3E-50 | 2E-48 | Tnfsf11 (TRANCE) | 3.2 | 6.6 | 67 | 3E-16 | 4E-14 |
| **Csf2** | 11 | 4.8 | 208 | 4E-47 | 3E-45 | Il15 | 1.9 | 0.88 | 6.7 | 0.0096 | 0.03 |
| **Il17f** | 11 | 4.9 | 171 | 5E-39 | 2E-37 | Tgfb3 | 1.4 | 4.0 | 16 | 6E-05 | 0.0005 |
| **Lif** | 10 | 3.6 | 132 | 1E-30 | 5E-29 | Il6st | 1.3 | 6.7 | 9.9 | 0.002 | 0.008 |
| Il22 | 9.1 | 2.4 | 43 | 4E-11 | 4E-10 | Il17f | 1.1 | 4.9 | 8.5 | 0.004 | 0.02 |
| Il21 | 7.5 | 0.65 | 23 | 2-06 | 1E-05 | Il21 | -3.8 | 0.65 | 11 | 0.0008 | 0.005 |
| **Tnf** | 4.5 | 6.0 | 188 | 1E-42 | 6E-41 | Il17a | -2.4 | 7.3 | 38 | 9E-10 | 38E-08 |
| **Lta (TNFb / lymphotoxin A)** | 3.8 | 6.2 | 110 | 9E-26 | 3E-24 | Ifng | -2.4 | 5.9 | 35 | 3E-09 | 8E-08 |
| **Il1b** | 2.6 | 4.5 | 6.2 | 0.01 | 0.03 | Tnfsf10 (TRAIL) | -1.6 | 4.0 | 19 | 2E-05 | 0.0002 |
| Tnfsf11 (TRANCE) | 2.3 | 6.6 | 35 | 3E-09 | 2E-08 | Lif | -1.4 | 3.6 | 11 | 0.0009 | 0.005 |
| **Csf1 (M-CSF)** | 2.3 | 3.0 | 11 | 0.0009 | 0.003 |  |  |  |  |  |  |
| Tnfsf10 (TRAIL) | 1.1 | 4.0 | 9.7 | 0.002 | 0.005 |  |  |  |  |  |  |

| Reinfection MAIT cells |  |  |  |  |  |
| --- | --- | --- | --- | --- | --- |
| v |  |  |  |  |  |
| Acute infection MAIT cells |  |  |  |  |  |
| Gene | Log fold change | Log CPM | LR | P value | FDR P value |
| Csf1 (M-CSF) | 2.2 | 3.0 | 12 | 0.0006 | 0.01 |
| Il6st | 1.5 | 6.7 | 14 | 0.0002 | 0.005 |
| Tnfsf11 (TRANCE) | 1.3 | 6.6 | 14 | 0.0002 | 0.007 |
| Il17a | -1.5 | 7.3 | 15 | 9.7E-05 | 0.003 |
| Csf2 (GM-CSF) | -1.2 | 4.8 | 12 | 0.0006 | 0.01 |

Genes shown are censored at FDR $P \leq 0.05$ and log(2) fold change of $\pm 1$ , and ordered by log fold change. Genes highlighted in bold are
differentially expressed in both human and mouse. CPM, Counts per million; FDR, false discovery rate; LR, likelihood ratio.

Similar analysis of chemokines showed strong activation-induced upregulation of a range of chemokines, including XCL1, CCL3 (MIP1α), CCL4 (MIP1β), and CXCL16 common to both species (Supplementary tables S10, S11), and of a common array of chemokine receptors CCR6, CXCR6, CCR1, CCR2 and CCR5 (Supplementary tables S12,S13, Supplementary figure S2), underlining marked evolutionary conservation of MAIT cell function.

### Pathway analysis of the MAIT cell transcriptome

To analyse the transcriptome at the level of pathways, rather than individual genes, we looked for upregulation of pathways using the open source, manually curated, peer-reviewed Reactome database^16^. The main pathways upregulated in human 5-OP-RU stimulated MAIT cells compared with naïve CD8^+^CD45RA^+^ T cells were related to endoplasmic reticulum stress – the unfolded protein response, and the related pathways IRE-1-a activation of chaperones and XBP1(S) activation of chaperones – to chemokine receptor-ligation, and to cholesterol biosynthesis (Figure 2A). When stimulated human MAIT cells were contrasted directly with unstimulated MAIT cells the activation of chemokine and cytokine signalling pathways – chemokine receptor-ligation, IL-2 signalling, interleukin receptor SHC signalling – was more apparent, as was human solute carrier-mediated transmembrane transport (Figure 2B).

**Figure 2.**
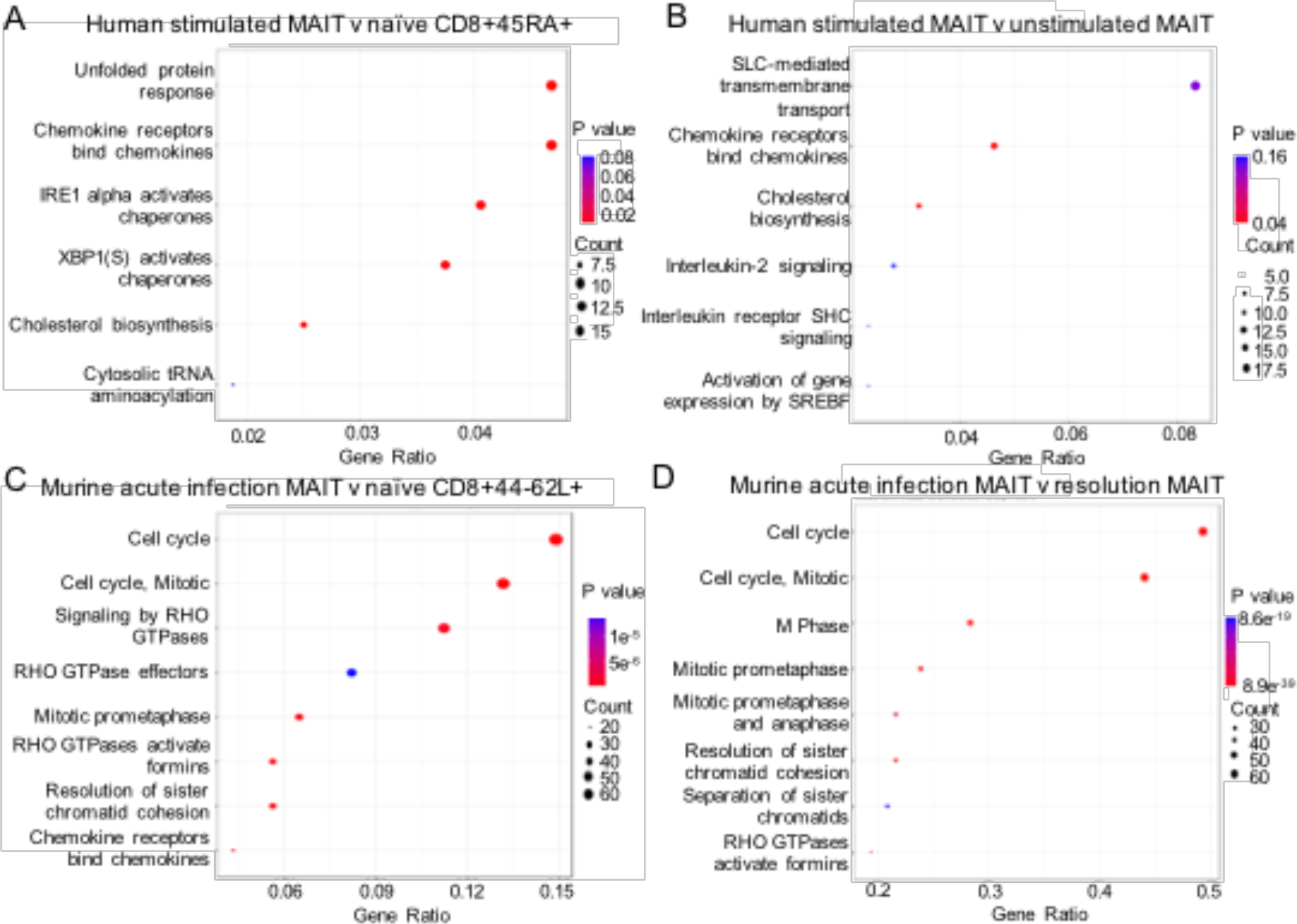
Reactome pathway analysis of activated MAIT cells. Pathway analysis of human and murine activated MAIT cell transcriptomes. Human peripheral blood 5-OP-RU-stimulated MR1-5-OP-RU-tetramer^+^ MAIT cells compared with **(A)** naïve CD8^+^CD45RA^+^ cells or **(B)** unstimulated MAIT cells. Murine pulmonary MR1-5-OP-RU-tetramer^+^ MAIT cells day 7 post-infection with *Legionella* were compared with **(C)** naïve CD8^+^CD44^-^CD62L^+^ T cells from uninfected mice or **(D)** MR1-tetramer^+^ MAIT cells from mice 12 weeks post infection with *Legionella*. Plots show the extent to which named pathways from the curated Reactome database are upregulated. Colourintensity represents statistical significance of the upregulation, dot size represents the number of genes upregulated in the pathway, x axis represents the proportion of all differentially expressed genes included in the pathway (‘gene ratio’). Pathways were selected using a significance threshold of a log fold change > 2 and P <0.01.

Perhaps reflective of the different context of activation, and consistent with the very rapid MAIT cell expansion observed with infection *in* vivo^10^, the murine MAIT cells activated by acute *L. longbeachae* infection showed very strong activation of cell cycle pathways as well as signalling by RHO GTPases and chemokine receptor-ligation, with similar dominance of the cell cycle when MAIT cells activated by acute infection were contrasted directly with unstimulated MAIT cells after infection resolution (Figure 2C,D).

### Comparison of MAIT cell transcriptomic profile with other T cell subsets

MAIT cells are a relatively ancient T cell subset, with both innate and adaptive properties, and are capable of expressing diverse functions depending on the nature of the pathogenic encounter^10,17^. Therefore we sought next to explore the nature of the murine MAIT cell transcriptome by comparing it with the transcriptional profiles for a wide range of other cell types reported within the Immunological Genome Project database^18^. Using hierarchical clustering, whilst the pulmonary naïve CD8^+^CD44^-^CD62^+^ T cells clustered with reference naïve CD8^+^ splenic T cells, activated MAIT cells from acute primary infection or from acute reinfection clustered most closely to invariant NKT cells (Figure 3). By contrast, after resolution of the infection MAIT cells clustered most closely with unstimulated splenic γδ T cells.

**Figure 3.**
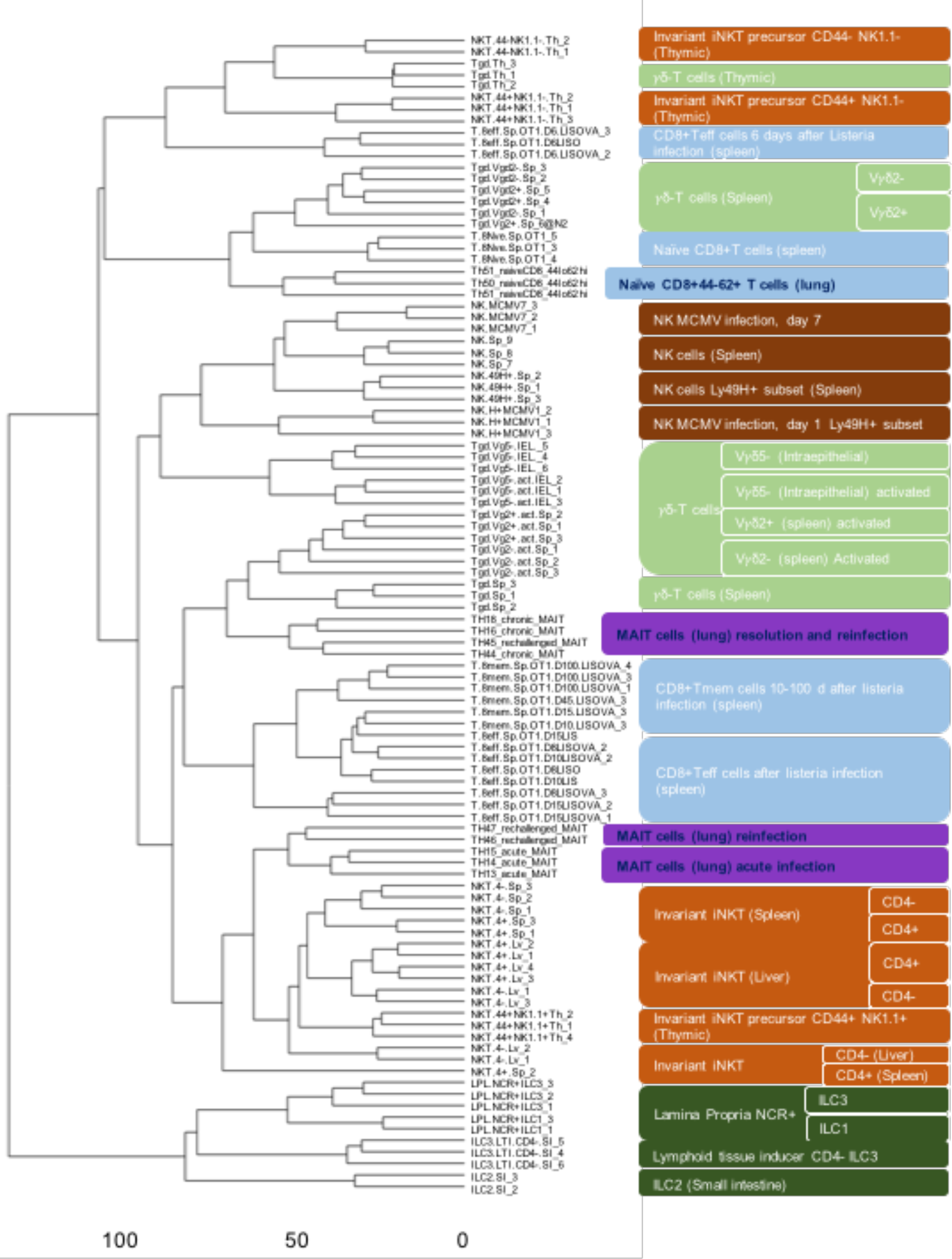
Comparison of murine MAIT cell transcriptomes with other cells in Immunological Genome Project dataset. Hierarchical clustering was used to compare transcriptomes of murine pulmonary MAIT cells or naïve CD8^+^CD44^-^CD62L^+^ cells with eighty-eight other cell types deposited in the Immunological Genome Project (ImmGen) database. Figure shows a dendrogram (left), ImmGen identifiers (middle) and the full name of each cell type (right). ImmGen samples are identified in white lettering. Samples from the current study are identified in black lettering, with extended lozenges. Cell types are colour-coded: invariant natural killer T cells (iNKT, orange), natural killer (NK) cells (brown), γδ T cells (light green), innate lymphoid cells (ILC, dark green), conventional CD8 T cells (blue), MAIT cells (purple). CD, clonal designation; NCR, NK cell receptor; Teff, effector T cell; Tmem, memory T cell.

### MAIT cells express a tissue repair transcriptional profile

Our observation of a distinct cytokine signature after infection resolution suggested that MAIT cells might be capable of performing more diverse functions than a purely pro-inflammatory response to TCR ligation. As observed already, TCR ligation in the absence of a TLR-agonist did not induce proliferation of murine MAIT cells. A wide variety of bacteria, mycobacteria and yeasts, including many commensal organisms^19^ express the riboflavin biosynthetic pathway, and may therefore be a major source of activating MR1 ligands, constitutively, or during breach of a barrier surface. Indeed MAIT cells require commensal organisms for their expansion^20^. A class of skin-homing Tc17 cells specific to commensal flora has recently been described, which expresses a ‘tissue repair’ gene signature and can accelerate repair of an epithelial wound^21^. These cells share several features with non-classical T cells, including the Type-17 cytokine profile and restriction by another MHC class 1b antigen presentation molecule H2-M3. Therefore we asked whether this tissue repair phenotype was a shared transcriptional programme in MAIT cells. We used gene set enrichment analysis^22^ (GSEA) to compare expression of this set of tissue repair genes^21^ (Supplementary table S14) with genes differentially expressed in MAIT cells. Indeed this gene set was markedly enriched in human MAIT cells after 5-OP-RU stimulation (normalised enrichment score (NES) 1.38, family-wise error rate (FWER) P<0.01, Figure 4A, B, Table 5). Similarly, despite differences in species, timecourse and method of MAIT cell activation, the same gene set was even more highly enriched in mice during acute *L. longbeachae* infection (NES 1.38, FWER p<0.01, Figure 4C, D, Table 5), with enrichment of ten genes common to both analyses (TNF, CSF2, HIF1A, FURIN, VEGFB, PTGES2, PDGFB, TGFB1, MMP25).

**Table 5.**
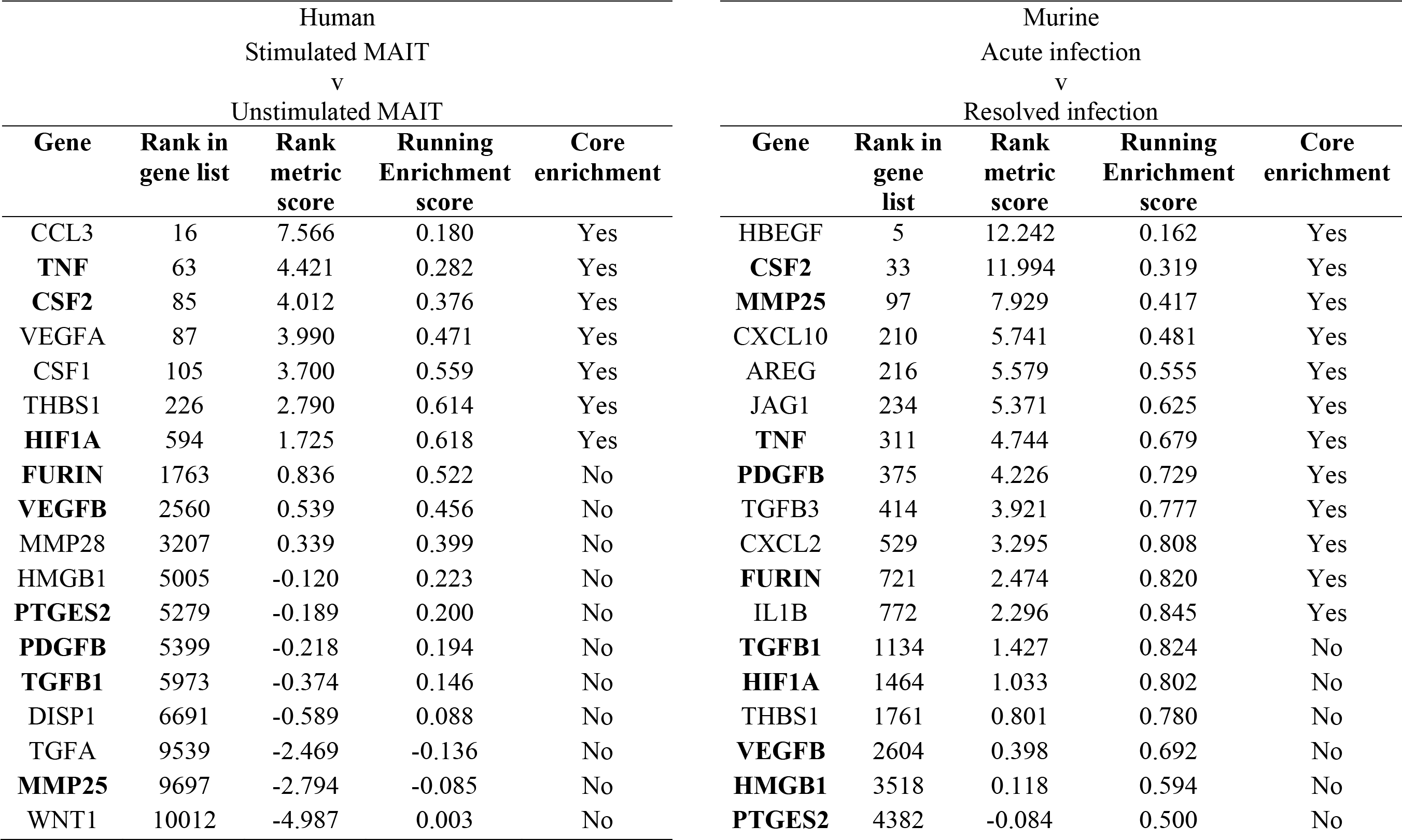
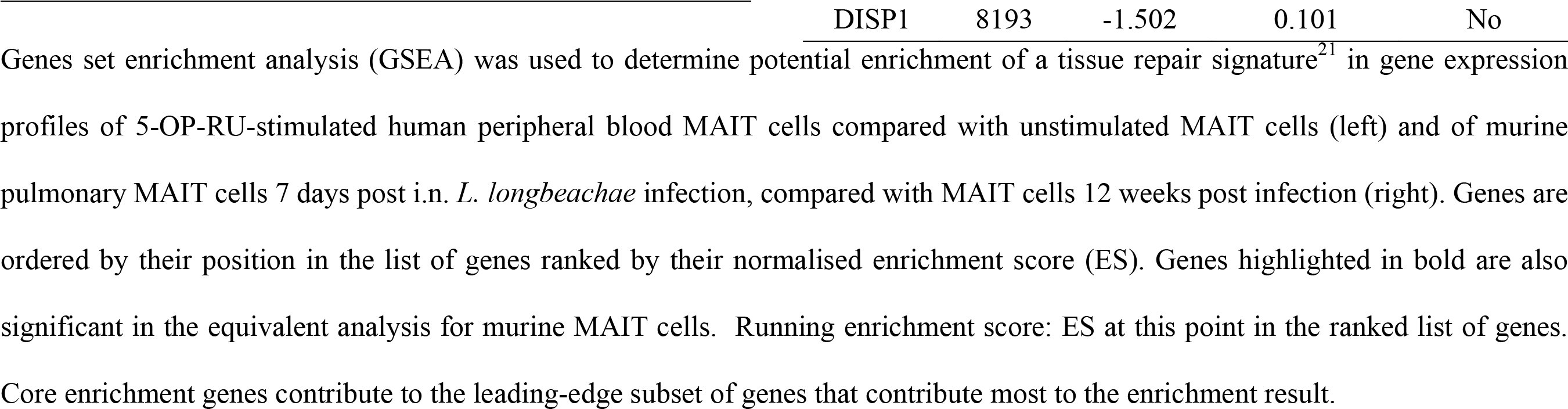
Gene set enrichment analysis for Tissue Repair set.

**Figure 4.**
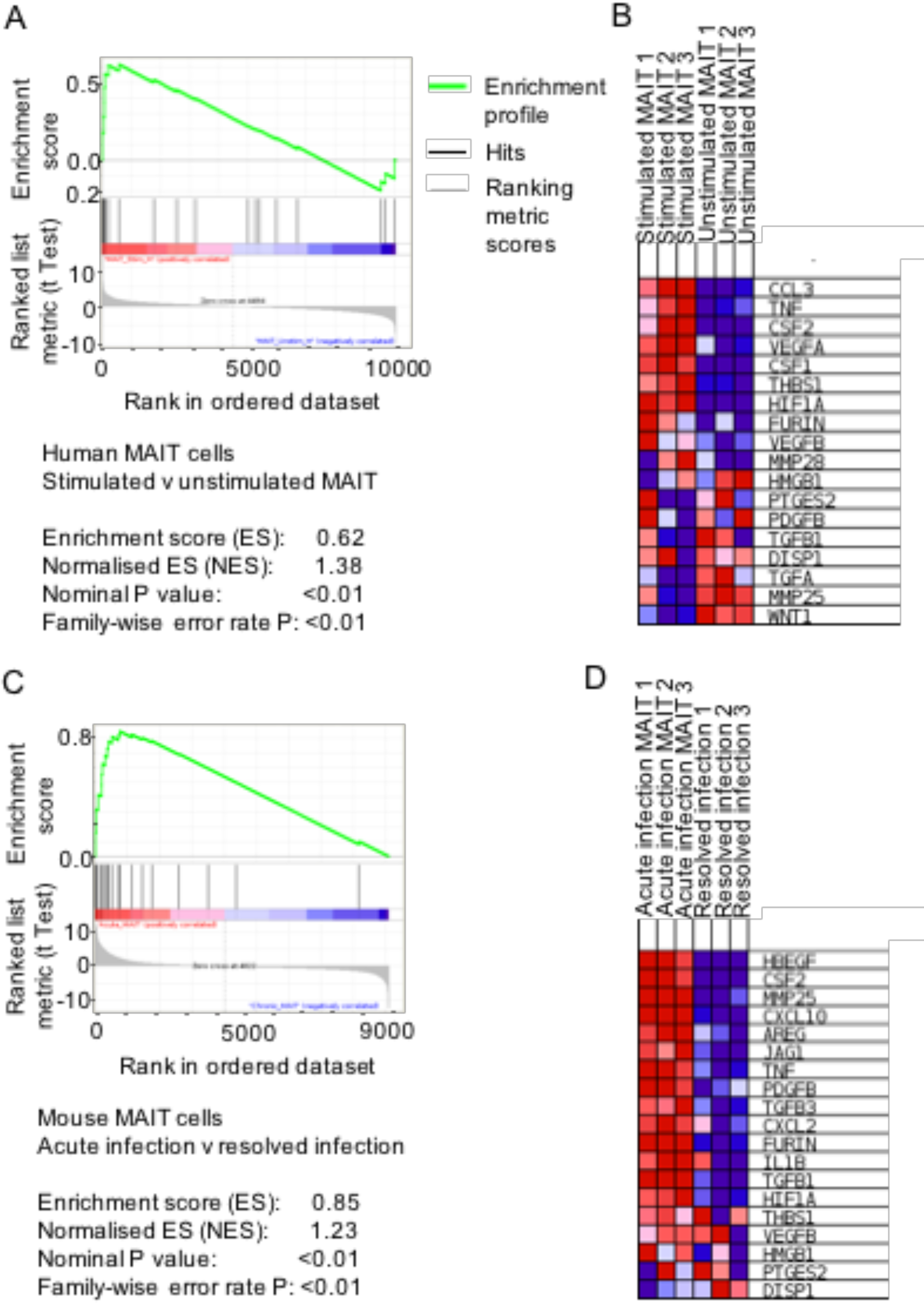
Gene set enrichment analysis for tissue repair gene signature in human and murine MAIT cells. Gene set enrichment analysis (GSEA) was used to determine potential enrichment of a tissue repair signature^21^ in gene expression profiles from human (A,B) and murine (C,D) MAIT cells. **(A)** GSEA summary plots for 5-OP-RU-stimulated human peripheral blood MAIT cells compared with unstimulated MAIT cells. The gene set is highly enriched:enrichment score (ES) = 0.62, normalised enrichment score (NES) = 1.38, nominal p value <0.01, family-wise error rate (FWER) p value <0.01. **(B)** Heat map of expression of leading edge subset genes within the gene set (red, highest expression, blue, lowest). **(C)** GSEA summary plots for murine pulmonary MAIT cells 7 days post i.n. *L. longbeachae* infection, compared with MAIT cells 12 weeks post infection. The gene set is highly enriched: enrichment score (ES) = 0.85, normalised enrichment score (NES) = 1.23, nominal p value <0.01, family-wise error rate (FWER) p value <0.01. **(D)** Heat map of expression of leading edge subset genes within the gene set (red, highest expression, blue, lowest).

## Discussion

Here we have systematically investigated the requirements for TCR-mediated activation of MAIT cells in mice, and delineated both *ex vivo* in human and *in vivo* in mice the consequences of this activation at a transcriptomic level. Due to their pro-inflammatory cytokine profile^8^ and specificity for a restricted selection of microbially-derived small molecules^2,23^, the most immediately apparent function of MAIT cells has hitherto been the early detection of microbes and initiation of an inflammatory host response^7, 11, 12, 24, 25^. Consistent with previous studies, our data confirm MAIT cells’ capacity for a strong, rapid pro-inflammatory response. However, in contrast to the similarities between activated MAIT cells and iNKT cells, the close similarity at a transcriptional level of resting murine MAIT cells to γδ T cells and the discovery of a clear transcriptional signature for tissue repair suggests that MAIT cells potentially have much broader roles in mucosal immunity.

The nature of these roles may depend on the context and nature of the cells’ activation. As with iNKT cells^26^ MAIT cells may be activated either via TCR recognition of ligand presented on MR1, or via cytokines alone, in the absence of a TCR signal, as occurs during respiratory viral infection ^17,27,28^. As we have observed previously, in the absence of inflammatory cytokines TCR-ligation alone is not sufficient to produce MAIT cell proliferation and activation *in vivo*^9^. Rather a second signal is required. *In vitro* in humans it has been shown that agonists of TLR1, TLR2, and TLR6 can provide this costimulus to drive MAIT cell cytokine secretion^29^. Consistent with, and extending these previous observations^9^, we observed here that murine MAIT cells proliferated in response to a different, but similarly restricted, set of TLR agonists: those for TLR3, TLR4, TLR6/2 and TLR9 and, but not for other TLRs tested.

As MAIT cells are found at their highest frequencies in the lungs and liver^11,30,31^ and the MR1-axis is highly conserved^5^ – implying a strong evolutionary pressure – it could be expected that potent MAIT cell responses should be elicited by a range of major human pathogens, and yet to date a strong human clinical phenotype for MAIT cell deficiency has not been described, and even in animal models the protective effect afforded by MAIT cells against mortality has been relatively modest in fully immune competent hosts^10,11^, likely due to multiple layers of immunological redundancy provided by other arms of innate and adaptive immunity^10^. One explanation might be that a different, more subtle function of MAIT cells explains Nature’s ongoing requirement for this subset in effective mucosal immunity.

Our data suggest an entirely novel function for MAIT cells in tissue repair. In both human and murine datasets, activated MAIT cells highly express a shared a gene expression signature with murine H2-M3 restricted commensal-specific Tc17 cells recently reported by Linehan *et al*^21^. Using topical skin colonisation with a specific clade of *S. epidermidis* this group demonstrated that specific commensal-derived *N*-formylated peptides presented on H2-M3, another class 1b MHC molecule, could induce tissue-resident Tc17 cells which provided specific capacity to promote tissue repair and remodelling (MMP25, Furin, PDGFB, TGFB1) and angiogenesis (CSF2, VEGFA, PDGFB)^21^. Healing of skin wounds was shown to be accelerated by colonisation of H2-M3 sufficient mice with these commensals. A human equivalent of H2-M3 has yet to be identified, but MAIT cells are abundant at barrier sites, allowing close interactions with commensal bacteria possessing an intact riboflavin metabolic pathway. Similarly to H2-M3 restricted CD8^+^ T cells, in this position MAIT cells are poised to maintain tissue homeostasis in the presence of commensals thereby limiting inflammation and associated tissue injury^32^. These data are consistent with a similar finding of this same tissue repair signature observed by GSEA analysis of MAIT cells when activated by TCR triggering, but not observed in the context of cytokine-mediated activation^33^. These findings might also explain the increased gut permeability observed in *Mr1*^-/-^ NOD mice compared with *Mr1*^+/-^ NOD littermates, which suggested a protective role for MAIT cells for maintaining gut homeostasis^34^. It was been speculated this might be mediated by IL-17A and IL-22^34^ which are both important in intestinal homeostasis^35,36^, and in the case of IL-22 induction of protective mucus-producing goblet cells^37^. In our dataset both cytokines were strongly upregulated in activated murine MAIT cells. Thus during mucosal damage, riboflavin-synthesising pathogens and commensal organisms might provide the MAIT cell activation both to induce the necessary inflammatory response to ensure bacterial clearance and also the signals necessary to accelerate healing of the wound. After successful clearance of infection, or barrier repair, the subsequent reduction in MR1-Ag presentation would ensure this signal declined.

Commensals might drive the MR1-MAIT cell axis in other ways. MAIT cell expansion requires exposure to a commensal microbiome^20^. Furthermore, commensal microbes have been implicated in enhancing host immunity against pathogens in the respiratory tract. In mice a Nod2-mediated IL-17A response to upper respiratory tract commensals enhanced CSF2 (GM-CSF) to promote bacterial killing and clearance by alveolar macrophages^38^. The strong upregulation of CSF2 (GM-CSF) we observed following TCR stimulation in human and murine MAIT cells would be beneficial in clearance of pathogenic microorganisms which have crossed the mucosal barrier. During tissue homeostasis commensal-derived MR1 signals might drive lower level, constitutive expression of GM-CSF needed to maintain alveolar macrophages in a pathogen responsive state^38^.

Another novel, prominent feature of MAIT cell activation in both humans and mice was the marked expression of the IL-6 family cytokine leukaemia inhibitory factor (LIF). Consistent with our finding of a MAIT cell tissue-repair signature, LIF has been found to be protective against epithelial damage in murine models of pneumonia^39^. LIF is significantly induced during pneumonia and can reduce lung epithelial cell death, promoting the expression of tissue-protective genes essential to lung regeneration and repair, and increased mucosal barrier integrity.

A unique feature of our transcriptomic dataset is that in a single experiment we were able to analyse MAIT cells from two different species obtained from two different tissues, using different contexts of activation, and yet we observed that the transcriptomic profiles of these MAIT cells were in fact very similar. Thus the distinctive, common properties of MAIT cells predominate over differences between these cells which might be observed in different contexts. Again, this underlines a strong conservation of functions likely driven by a consistent role in mucosal immunology.

Given the wide diversity of conventional and non-classical T cells now recognised^24^, many of which share common transcriptional programmes^40,41^ we applied a comparative approach^42^ to analyse the phenotype of MAIT cells, overcoming significant methodological hurdles to compare our RNA sequencing data directly with older microarray expression data in the ImmGen dataset. Activated MAIT cells were most similar to activated invariant iNKT cells, as might be expected from the similarities in surface markers and functional phenotype^5,24,43^. This is likely related to shared transcriptional signatures controlled by common transcription factors, not least that which has been described for promyelocytic leukemia zinc finger (PLZF) which defines a distinct surface phenotype and functional capacity in CD161^+^ NK cells, iNKT cells and MAIT cells^40^. However it is interesting that in their resting state MAIT cells more closely resembled splenic γδ T cells. Unlike MAIT and iNKT cells, most γδ T cells are not constrained by a specific MHC restriction^44,45^, rather they have different functional profiles associated with usage of different TCR V gene segments. Depending on the Vγ subset, γδ T cells recognise a diverse range of small microbial metabolites, lipids, self-antigens and stress-induced proteins, and may display a range of functions associated with variously with inflammation, immunoregulation, cytotoxicity, antigen presentation^24^ and promotion of tissue repair^46^. In the absence of the TCR/TLR mediated activation, MAIT cells may be fulfilling a different, perhaps homeostatic function. Indeed, we were able to investigate what this might be by analysing the transcriptome of MAIT cells in their resting state, outside the context of inflammation. After resolution of infection, Tnfsf18 (Glucocorticoid-Induced TNF-Related Ligand, GITRL) is the most strongly upregulated cytokine. The function of GITRL is context dependent, but under resting, non-inflammatory it can negatively regulate NK cells and maintain or expand regulatory T cells conditions^47^. Other immunoregulatory cytokines were also upregulated: Tnfsf11 (TRANCE, RANKL) was identified in a commensal-derived immunoregulatory signature^21^, whilst IL-15 can inhibit T cell apoptosis to maintain memory T cell survival^48^. Together these data implicate resting MAIT cells in potentially significant immunoregulatory roles.

In summary, our analysis of TCR-activated MAIT cells demonstrates pronounced conservation of functions and gene expression profiles between human and murine cells, and suggests that beyond type 17 / type 1 pro-inflammatory responses to invading microbial pathogens, MAIT cells have the capacity to contribute to immunoregulatory and tissue repair roles likely to be essential for maintaining the integrity of mucosal barrier surfaces in health and disease.

## Materials and Methods

### Animal models

C57BL/6 mice were bred and housed in the Biological Research Facility of the Peter Doherty Institute (Melbourne, Victoria, Australia). Mice aged 6-12 weeks were used in experiments, after approval by the University of Melbourne Animal Ethics Committee (1513661 and 1513712).

Intranasal (i.n.) inoculation with a stimulatory MR1 ligand (76 pmol 5-OP-RU) or a non-activating MR1 ligand (76 pmol 6-FP) and TLR agonist (see Table S1) in a total 50 μl volume was performed on isofluorane-anaesthetised mice on day 0. Additional doses of the relevant MR1 ligand in 50 μl were administered on days 1, 2, and 4. Control mice received TLR agonists alone, in a total volume of 50 μl. For infection experiments mice were inoculated with 1-2 x10^4^ CFU *Legionella longbeachae* (clinical isolate NSW150) in 50 μl PBS.

Mice were weighed daily and assessed visually for signs of disease, including inactivity, ruffled fur, laboured breathing, and huddling behaviour. Animals that had lost ≥15% of their original body weight and/or displayed evidence of pneumonia were euthanised.

Mice were killed by CO_2_ asphyxia, the heart perfused with 10 ml cold Roswell Park Memorial Media-1640 (RPMI, Gibco) and lungs were taken. To prepare single-cell suspensions lungs were finely chopped with a scalpel blade and treated with 3 mg.ml^-1^ collagenase III (Worthington, Lakewood, NJ), 5 μg/ml DNAse, and 2% foetal calf serum in RPMI for 90 min at 37°C with gentle shaking, and, where relevant, brefeldin A (GolgiPlug™, BD Biosciences, San Diego, CA). Lung cells were then filtered (70 μm) and washed with PBS/2% foetal calf serum. Red blood cells were lysed with hypotonic buffer TAC (Tris-based amino chloride) for 5 min at 37°C. Approximately 1.5x10^6^ cells were filtered (40 μm) and used for flow cytometric analysis. To obtain sufficient cells for sorting naïve CD8^+^CD44^-^CD62L^+^ cells from infection-naïve mice lungs from 2-3 mice per sample were pooled and stained with 0.18 μl anti-CD8-PE, then magnetically enriched using anti-PE beads (Miltenyi) prior to sorting.

For analysis of systemic MAIT cell distribution lymphocytes were obtained from mesenteric lymph nodes by passing through a 70 μm strainer. Splenocytes were obtained by homogenising splenic tissue through a 70 μm strainer then preforming red cell lysis prior to staining. Peripheral blood cells were obtained from the inferior vena cava into a heparinised syringe and underwent surface staining prior to red cell lysis with 1 ml of 10%red cell lysis buffer (BD Bioscience) for 5 minutes at room temperature before washing twice with FACS buffer. Hepatic lymphocytes were obtained by perfusing the liver with 8-10 ml PBS, passing through a 40 μm strainer, washing once with PBS the resuspending in 36% Percoll (Sigma) and centrifuging without braking at 800 *g* for 25 mins at RT over a 70% Percoll underlay. Cells from the interphase were washed with FACS buffer, red cells lysed with 1 ml 10% red cell lysis buffer, cells washed twice and stained for flow cytometry.

### Determination of bacterial counts in infected lungs

Bacterial infection was determined for *L. longbeachae* by counting colony-forming units (CFU) obtained from plating homogenised lungs in duplicate from infected mice (x5 per group) on buffered charcoal yeast extract agar (BYCE) containing 30 μg/ml streptomycin and colonies counted after 4 days at 37°C under aerobic conditions. Culture media for other bacteria are shown in table S2.

### Antibodies flow cytometry and cell sorting

Details of flow cytometry antibodies are shown in Supplementary table S15. To block non-specific staining, cells were incubated with MR1-6-FP tetramer and anti-Fc receptor (2.4G2) for 15 min at room temperature and then incubated at room temperature with Ab/tetramer cocktails in PBS/2% foetal calf serum. Dead cells were excluded using 4′,6-diamidino-2-phenylindole (DAPI) for live cell sorting added for 10 mins or by staining for 20 mins in PBS with fixable viability dyes Zombie Yellow (Biolegend, 1:100, 423104) or Live/Dead EF780 (BD Bioscience, 1:1000, 565388).

For live cell sorting on human peripheral blood mononuclear cells (PBMC) 50ml of heparinised blood were obtained freshly per volunteer, mixed with an equal volume of phosphate buffered saline (PBS) and layered over an equal volume of Ficoll-Paque (GE Healthcare, Chicago, IL) and centrifuged at 800 *g* for 20 mins at room temperature. Cells were washed twice with PBS, cells counted by trypan blue estimation, then half the cells were resuspended overnight in RPMI with 10% human serum for overnight rest and the other half resuspended in flow cytometry buffer comprising PBS with 2% foetal calf serum and 2 mM EDTA (FACS buffer) for immediate magnetic enrichment and sorting. These cells were stained with surface antibodies (CD3-PE-CF594, CD8-PerCPCy5.5, CD45RA-FITC, TCR Vα7.2-APC and FCγ block for 15 mins at RT, followed by staining with MR1-5-OP-RU tetramer-PE for 20 mins at RT. Tetramer positive cells were positively selected using anti-PE microbeads (10 μl per 10^-7^ cells, Miltenyi, Cologne, Germany) according to the manufacturer’s instructions. Cells were sorted immediately into ice cold PBS with 10% FCS using an FACSAria III cell sorter (BD Bioscience) selecting live CD3^+^TCR-Vα7.2^+^MR1-5OP-RU-tetramer^+^ MAIT cells from the positive fraction and live CD3^+^CD8^+^CD45RA^+^MR1-Tetramer^-^ cells from the negative fraction. The following day the remaining cells were stimulated for 6 h with 10 nM 5-OP-RU then magnetically enriched using MR1-5-OP-RU-tetramer-PE and anti-PE microbeads, and live CD3^+^TCR-Vα7.2^+^MR1-5-OP-RU-tetramer^+^ MAIT cells sorted in the same manner. Purity was checked and with an average of 98%. Immediately after sorting cells were centrifuged at 400 *g* for 5 mins then resuspended in 100 μL of RNA lysis buffer (Agilent Ltd, UK) with 0. 7 μL β-mercaptoethanol and stored at -80°C.

For live cell sorting of murine T cells, CD8 cells were magnetically enriched using anti-CD8-PE and anti-PE microbeads and live CD8^+^CD44^-^CD62L^+^ cells from uninfected mice, or live CD3^+^CD19^-^CD45.2^+^TCRβ^+^MR1-5-OP-RU-tetramer^+^ MAIT cells from previously infected mice were sorted as above.

For intracellular staining, cells were fixed with 1% paraformaldehyde prior to analysis on LSRII or LSR Fortessa or Canto II (BD Biosciences) flow cytometers. For intracellular cytokine staining Golgi plug (BD Biosciences) was used during all processing steps. Cells stimulated with PMA (phorbol 12-myristate 13-acetate;)/ionomycin (20 ng ml^-1^, 1μg ml^-1^, respectively) for 3 h at 37°C were included as positive controls. Surface staining was performed at 37°C, and cells were stained for intracellular cytokines using the BD Fixation/Permeabilization Kit (BD, Franklin Lakes, NJ) or transcription factors using the transcription buffer staining set (eBioscience) according to the manufacturers’ instructions.

For validation of key targets identified by RNA sequencing flow cytometry was performed on cryopreserved human PBMC from additional healthy human donors. Samples were defrosted into pre-warmed RPMI with 10% human serum, stained with anti-TCR-Vα7.2-PE or anti-TCR-Vα7.2-PE and magnetically enriched using anti-PE or anti-APC microbeads. 200,000 positively-selected TCR-Vα7.2^+^ cells or the negative fraction (for naïve CD8^+^45RA^+^ cells) were co-cultured for 5 hours in the presence of brefeldin A with 100,000 class I reduced (C1R) antigen presenting cells (APCs) which had been previously pulsed for 2 hours with 10 nM 5-OP-RU, or with naïve C1R cells (unstimulated control), or with PMA / ionomycin (20 ng ml^-1^, 1μg ml^-1^, respectively), or without any stimulation. Cells were then analysed by surface and intracellular cytokine staining as above. For validation of murine targets cells were isolated from uninfected mice or mice which had undergone intranasal infection 7 days prior (acute) or 12 weeks prior (resolution) or reinfection 7 days prior, and cytometrically analysed as described above.

### RNA sequencing

Cells were lysed in Agilent lysis buffer (Agilent Ltd., UK) containing 100 mM β-mercaptoethanol and passed through a QIAshredder device (Qiagen, Valencia, US), then RNA extracted using the Absolutely RNA Nanoprep Kit according to the manufacturer’s instructions, including using of DNase I. RNA libraries were prepared at the Melbourne Translational Genomics Platform, Department of Pathology (The University of Melbourne). Briefly, RNA quality and quantity were assessed using the Bioanalyzer 2100 RNA pico kit (Agilent technologies). The input total RNA was normalized to 250 pg per sample and median RIN was 9.7 (range 5.7-10.0). RNA was reverse transcribed and cDNA amplified by *in vitro* transcription with the SMART-Seq v4 Ultra Low Input RNA Kit for Sequencing (Clontech). First strand cDNA synthesis and tailing by reverse transcription was performed using Clontech’s proprietary SMART (Switching Mechanism at 5’ End of RNA Template) technology. Following first strand synthesis, cDNA was amplified 12 cycles by LD PCR using blocked PCR primers. Amplified cDNA was purified using AMPure XP prior to QC using the bioanalyser 2100 HS DNA kit (Agilent technologies). Library preparation of purified amplified cDNA was performed using Nextera XT library preparation (Illumina, AUS). Following QC, 150 pg of cDNA was tagmented (simultaneously fragmented with adaptors inserted) using Nextera transposons. Molecular barcodes were incorporated during 12 cycles of amplification followed by purification using AMPure XP. The libraries passed a quality checkpoint (Qubit and Bioanalyser HS DNA) prior to normalization and pooling before loading onto the HiSeq 2500 (Illumina, AUS) for paired end sequencing.

### Quality control and bioinformatics analysis of RNA sequencing data

RNA-seq reads were aligned to reference genome sequences using STAR^49^ aligner software. Mapped reads were assigned to genomic features using *Rsubread*^50^ R package^51^.

Genes that were differentially expressed (>2 fold, p<0.01, FDR<0.05) between conditions and their normalised expression values, were generated with *EdgeR* R package^52^, Partek^®^ Flow^®^, an online analysis platform for Next Generation Sequencing data (http://www.partek.com/partek-flow/). Pathway enrichment analysis using the Reactome platform^16^ was performed using *ReactomePA^53^* R package. Gene count data transform to log2-counts per million (logCPM) was performed using *voom* function in *limma*^*54*^ R package. Gene set enrichment analysis (GSEA) using *voom* transformed count data was performed using GSEA version 3.0^55^, comparing gene expression data as a whole with the reference gene list obtained from the publication by Linehan et al.^21^

RNA-seq data (logCPM) and ImmGen microarray data were integrated using a common set of *Entrez*^*56*^ annotated genes; batch effect removal was performed using *ComBat* algorithm in *sva*^*57*^ R package. Hierarchical clustering analysis of transcription profiles was conducted in R employing highly variable genes (IQR >0.75 ^58^) and Euclidian distance.

### Standard statistical analysis

Statistical tests were performed using the Prism GraphPad software (version 7.0 La Jolla, CA). Comparisons between groups were performed using Student’s t-tests or Mann-Whitney tests as appropriate unless otherwise stated. Flow cytometric data analysis was performed with FlowJo10 software (Ashland, OR).

### Reagents

Human peripheral blood mononuclear cells (PBMC) were obtained from the Australian Red Cross Blood Service (ARCBS) (University of Melbourne Human Research Ethics Committee 1239046.2). Healthy human lung explant tissue was obtained via the Alfred Lung Biobank program and ARCBS from organs not suitable for donation (Blood Service HREC 2014#14 and University of Melbourne Human Research Ethics Committee 1545566.1).

### Compounds, immunogens and tetramers

5-OP-RU was prepared as described previously^6^. CpG1668 (Sequence: T*c*C*A*T*G*A*C*G*T*T*C*C*T*G*A*T*G*C*T (*phosphorothioate linkage) nonmethylated cytosine-guanosine oligonucleotides was purchased from Geneworks (Thebarton, Australia) and Pam2Cys was chemically synthesised and functionally verified in house. Other toll like receptor ligands are detailed in Supplementary table S1. Murine and human MR1 and b2-Microglobulin genes were expressed in *Escherichia coli* inclusion bodies, refolded, and purified as described previously^59^. MR1-5-OP-RU tetramers were generated as described previously^2^.

### Bacterial strains

Cultures of *Legionella longbeachae* NSW 150 were grown at 37°C in buffered yeast extract (BYE) broth supplemented with 30-50 μg/ml streptomycin for 16 hours to log-phase (OD600 0.2-0.6) with shaking at 180 rpm. For the infecting inoculum, bacteria were re-inoculated in pre-warmed medium for a further 2-4 h culture (OD_600_ 0.2-0.6) with the estimation that 1 OD_600_=5x10^8^/ml, sufficient bacteria were washed and diluted in phosphate buffered saline (PBS) with 2% BYE for i.n. delivery to mice. A sample of inoculum was plated onto (BYCE) with streptomycin for verification of bacterial concentration by counting colony-forming units.

## Supporting information

## Acknowledgments

This work was funded by grants to T.S.C.H. from the Wellcome Trust (104553/z/14/z, 211050/Z/18/z). The research was supported by the National Institute for Health Research (NIHR) Oxford Biomedical Research Centre (BRC). The views expressed are those of the authors and not necessarily those of the NHS, the NIHR or the Department of Health. The work was also supported by the National Health and Medical Research Council of Australia (NHMRC) Program Grants 1113293, 1071916, 1016629 and 606788, and Project Grant 1120467. A.J.C. is supported by an ARC Future Fellowship. S.B.G.E. is supported by an ARC DECRA Fellowship. P.K. was supported by an NIHR Senior Fellowship, Oxford Martin School (PK) and the Wellcome Trust (WT109965MA). We are grateful to Dr Brendan Russ and Linda Wakim for assistance and suggestions for experimental design; Dr Ama Essilfie, Prof Richard Strugnell, Frances Oppodisam, Jennifer Davies, Prof Roy Robbins-Browne, Prof Kenneth Beagley and Hayley Newton for bacterial strains; Prof David Jackson for Pam2Cys; Dr Jeffrey Mak for MR1 ligands; Dr Vanta Jameson, Mr Josh Kie at the Flow Cytometry Facilities at the Melbourne Brain Centre and the Peter Doherty Institute; and to Kym Pham and Karey Cheong at the Melbourne Translational Genomic Platform.

## Author contributions

TSCH, TLP, LK, SBG, BSM, BR, KP, KC performed the experiments. TSCH, MO, EM, AK, MJ analysed the data. TSCH, JM, ZC, AC, ST, PK designed the experiments and managed the study. TSCH, MJ, AC, ZC, JM, PK conceived the work and wrote the manuscript which was revised and approved by all authors.

## Materials and Correspondence

Correspondence and material requests should be addressed to TSC Hinks.

---

**Supplementary figure S1.**
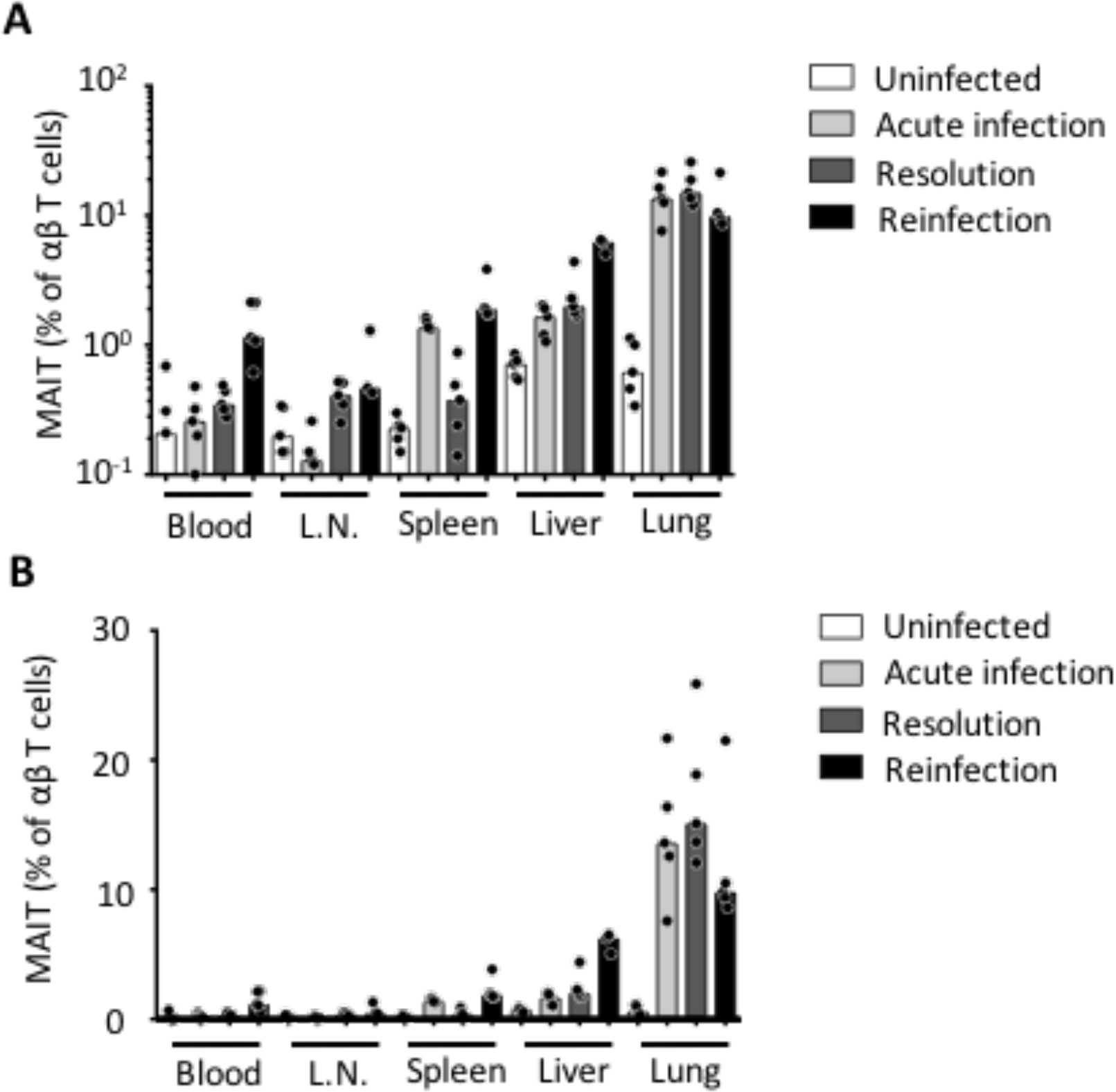
Tissue distribution of MAIT cells during infection *in vivo*. Relative frequencies of MR1-5-OP-RU tetramer^+^ MAIT cells as a proportion of total live TCRβ^+^ T cells in the peripheral blood, mesenteric lymph node (L.N.), spleen, liver and lungs of C57BL/6 mice before, or after intranasal infection with 1x10^4^ CFU *L. longbeachae*. Mice were sacrificed before (‘uninfected’), or 7 days after infection (‘acute’), or at least 12 weeks post infection (‘resolution’) or 7 days after a second intranasal infection with 2x104 CFU *L. longbeachae* in mice which had recovered from infection 12 weeks previously (‘reinfection’). Graph shows combined data from experiments using 3-5 mice per group and performed one-three times.

**Supplementary figure S2.**
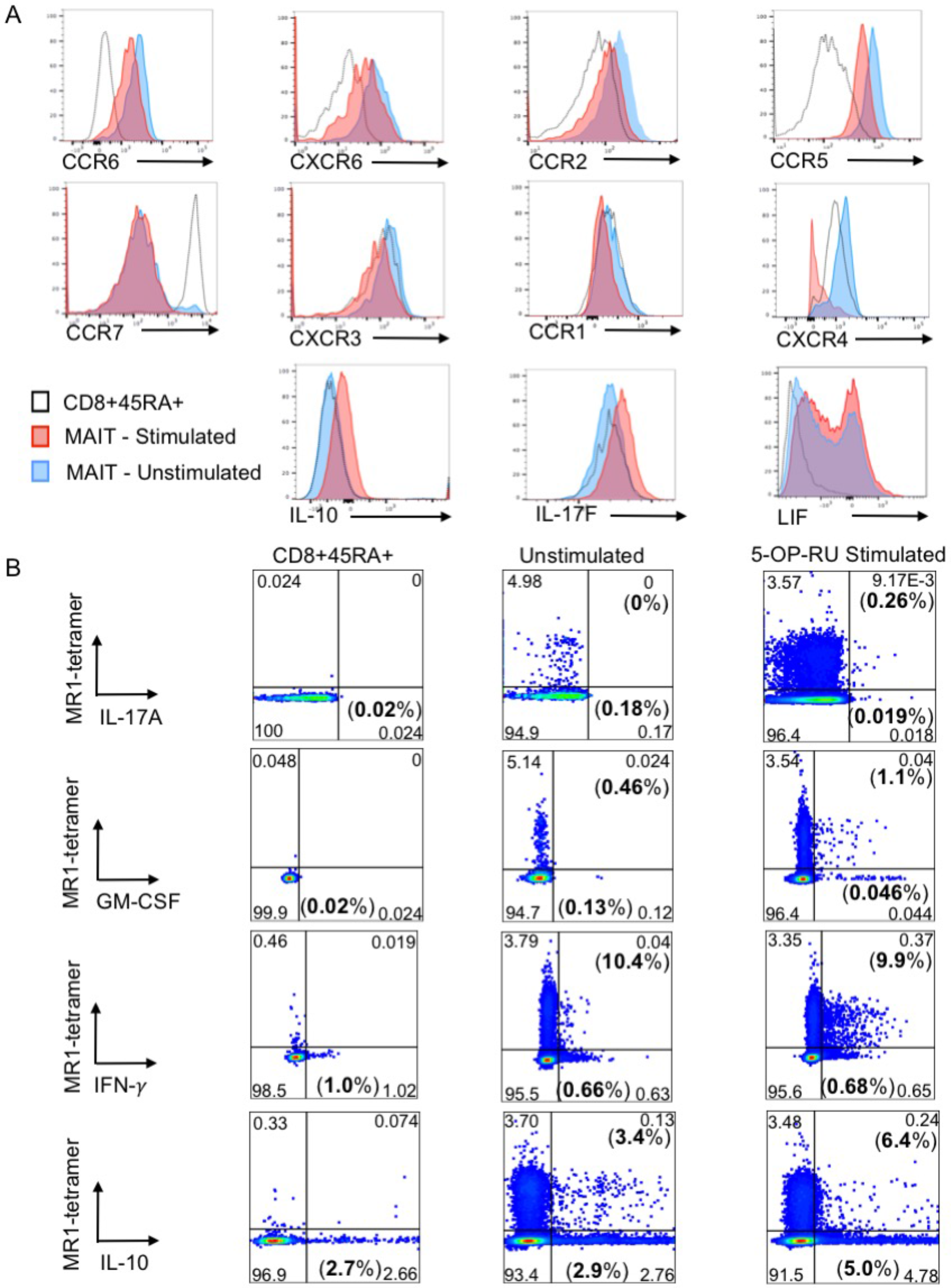
Cytometric validation of key differentially expressed genes (human) (**A**) Representative flow-cytometry plots showing surface expression of the chemokine receptors CCR5, CCR7, CCR2, CCR1, CXCR6, CXCR4, CCR6, CXCR3, and intracellular expression of the cytokines IL-10, IL-17F and LIF. Histograms compare staining of CD8^+^CD45RA^+^ cells (black, dotted) with unstimulated MAIT cells (blue) or MAIT cells after 6 h stimulation with 10 nM 5-OP-RU (red). (**B**) Representative flow-cytometry plots showing expression of the cytokines IL-17A, GM-CSF, IFN-γ and IL-10, by intracellular cytokine staining. Histograms compare staining of MR1-5-OP-RU tetramer^+^ MAIT cells after 6 h stimulation with 10 nM 5-OP-RU in the presence of brefeldin A (right) with unstimulated cells (middle) and tetramer-negative, CD8^+^CD45RA^+^ naïve T cells (left). Figures in brackets represent percentage of MAIT cells (top / middle) or of CD8^+^CD45RA^+^ T cells (bottom) expressing the cytokine. Results are representative of three independent donors.

**Supplementary figure S3.**
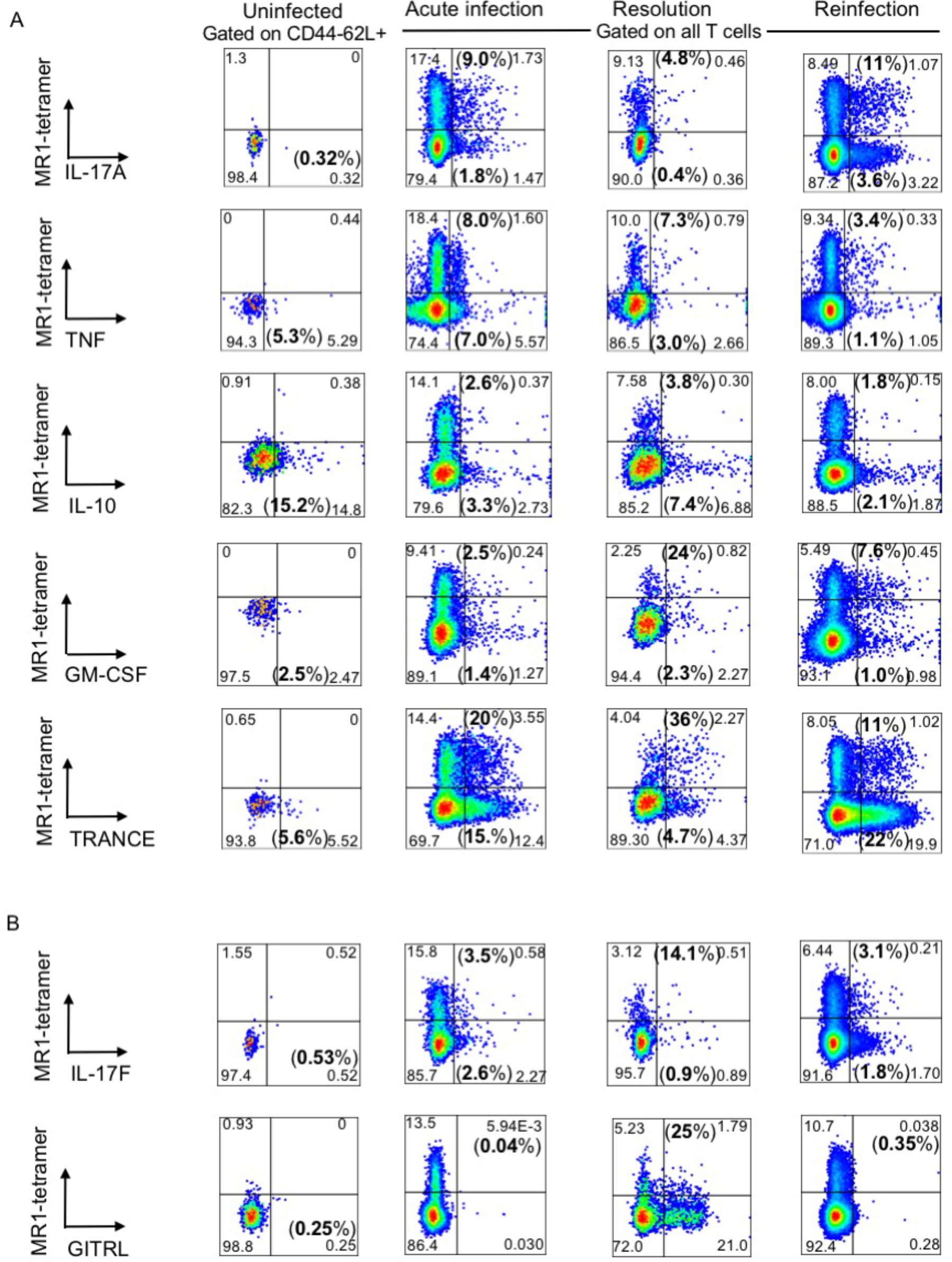
Cytometric validation of key differentially expressed genes (murine) Representative flow-cytometry plots showing expression of the cytokines IL-17A, TNF, IL-10, TRANCE (TNFSF11), GM-CSF, IL-17F and GITRL (TNFSF18) by intracellular cytokine staining on murine pulmonary T cells. Histograms compare staining of CD44^-^ CD62L^+^ T cells from uninfected mice (left) with MR1-5-OP-RU tetramer^+^ MAIT cells either 7 days (‘acute’, middle left) or 12 weeks (’resolution’, middle right) after infection with 1 x10^4^ CFU intranasal *L. longbeachae*, or 7 days after reinfection with 2 x10^4^ CFU i. n. *L. longbeachae* in mice previously infected 12 weeks prior with 10^4^ CFU i.n. *L. longbeachae* (‘reinfection’, right). Cells were incubated for 4 h in the presence of brefeldin A without (A, *ex vivo*) or with (B, stimulated) PMA and ionomycin. Figures in brackets represent percentage of CD44^-^CD62L^+^ T cells (left, lower quadrants) or MR1-tetramer^+^ MAIT cells (middle and right, upper quadrants), or MR1-tetramer^-^ conventional T cells (middle and right, lower quadrants) expressing the cytokine. Results are representative of three independent replicates performed on two separate days.

**Supplementary figure S4.**
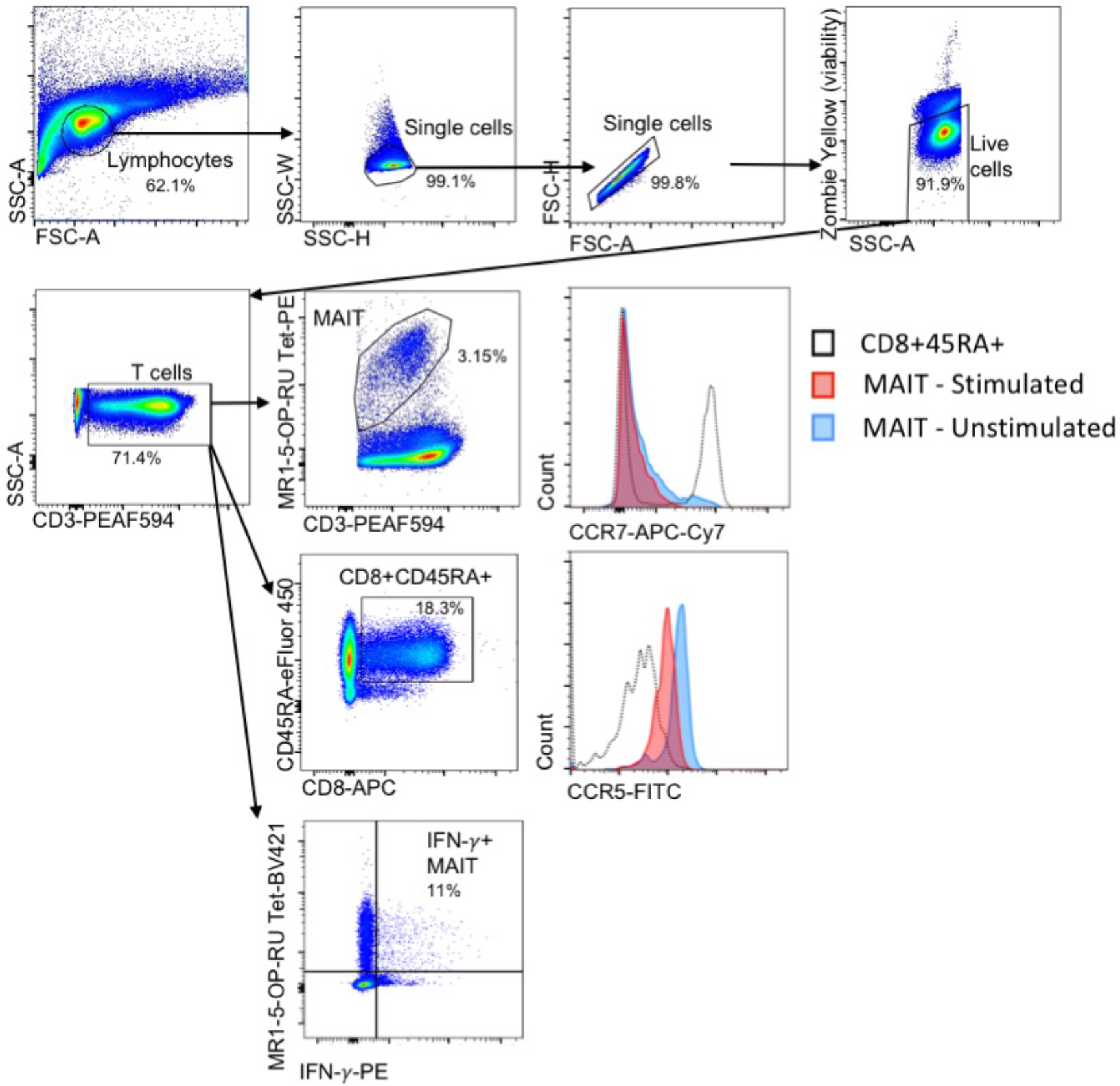
Cytometric gating strategy (human) Human peripheral blood lymphocytes were identified, and doublets excluded, using forward and side scatter characteristics. Dead cells were excluded using Zombie Yellow viability stain, then populations were gated on CD3^+^ cells, then on either MR1-5-OP-RU tetramer conjugated to PE or BV421 for MAIT cells, or on CD8 and CD45RA to identify naïve CD8^+^CD45RA^+^ cells. Histograms compare staining of CD8^+^CD45RA^+^ cells (black) with unstimulated MAIT cells (blue) or MAIT cells after 6 h stimulation with 10 nM 5-OP-RU (red). For intracellular cytokines, where basal cytokine secretion was minimal, gates were set on the unstimulated MAIT cell sample.

**Supplementary figure S5.**
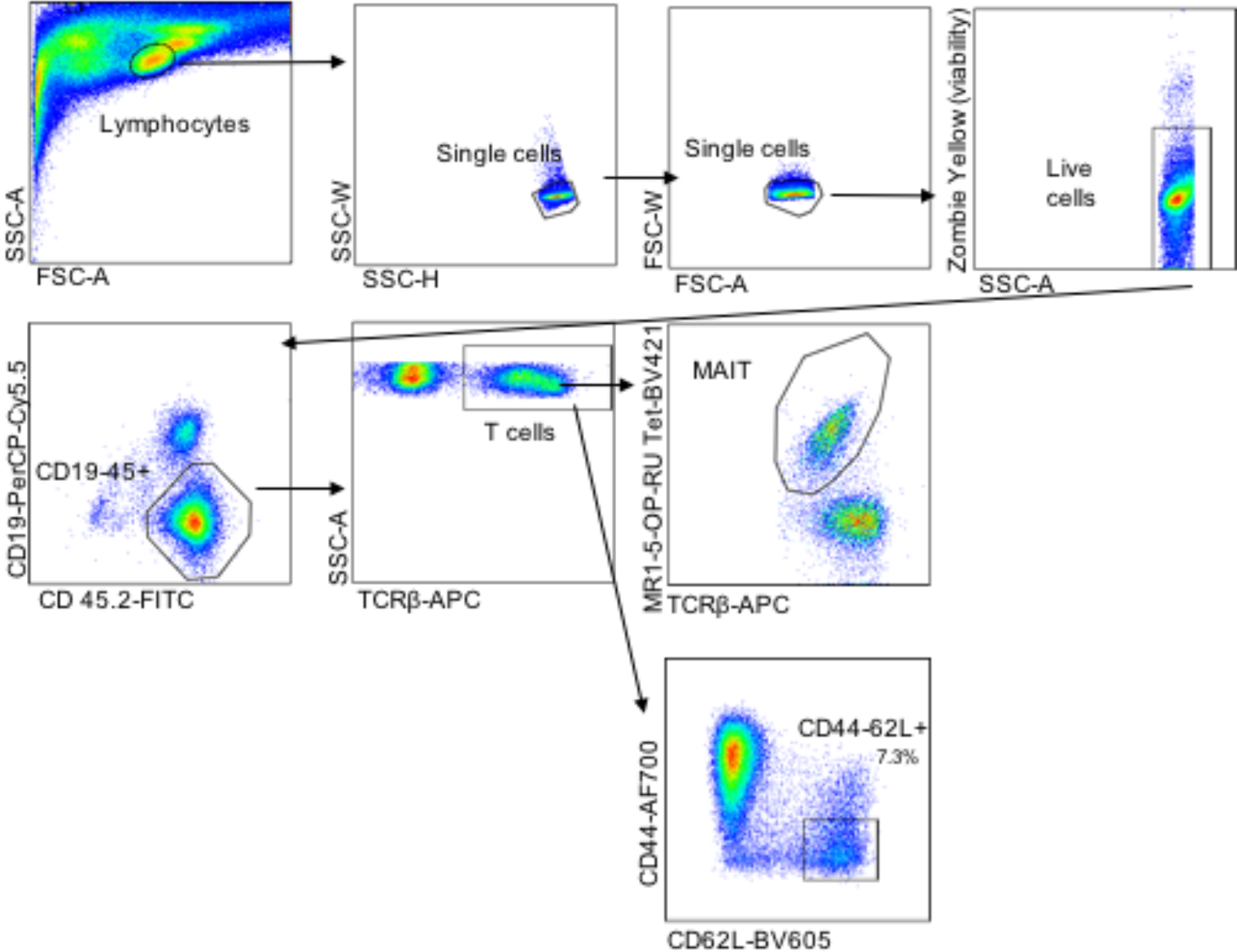
Cytometric gating strategy (murine) Pulmonary lymphocytes were identified, and doublets excluded, using forward and side scatter characteristics. Dead cells were excluded using Zombie Yellow viability stain, then populations were gated on CD19^-^CD45.2^+^ cells, then TCRβ^+^ cells and finally MR1-5-OP-RU tetramer^+^ MAIT cells, or CD44^-^CD62L^+^ T cells.

Supplementary table S1. Toll like receptor agonists used in murine experiments

Supplementary table S2. Comprehensive list of differentially expressed genes. Human.

Genes shown are censored at FDR P ≤0.05 and log(2) fold change of ±1, and ordered by log fold change. CPM, Counts per million; FDR, false discovery rate; LR, likelihood ratio.

Supplementary table S3. Comprehensive list of differentially expressed genes. Murine.

Genes shown are censored at FDR P ≤0.05 and log(2) fold change of ±1, and ordered by log fold change. CPM, Counts per million; FDR, false discovery rate; LR, likelihood ratio.

Supplementary table S4. TCR usage overrepresented genes. Human.

Genes shown are censored at FDR P ≤0.05 and log(2) fold change of ±1, and ordered by log fold change. CPM, Counts per million; FDR, false discovery rate; LR, likelihood ratio.

Supplementary table S5. TCR usage overrepresented genes. Murine.

Genes shown are censored at FDR P ≤0.05 and log(2) fold change of ±1, and ordered by log fold change. CPM, Counts per million; FDR, false discovery rate; LR, likelihood ratio.

Supplementary table S6. Differentially expressed cytokine receptors. Human.

Genes associated with formation of T cell memory which are found to be differentially expressed in this dataset. Genes shown are censored at FDR P ≤0.05 and ordered by log fold change. CPM, Counts per million; FDR, false discovery rate; LR, likelihood ratio.

Supplementary table S7. Differentially expressed cytokine receptors. Murine.

Genes shown are censored at FDR P ≤0.05 and log(2) fold change of ±1, and ordered by log fold change. CPM, Counts per million; FDR, false discovery rate; LR, likelihood ratio.

Supplementary table S8. Differentially expressed surface markers (Cluster of Differentiation molecules). Human

Genes shown are censored at FDR P ≤0.05 and log(2) fold change of ±1, and ordered by log fold change. CPM, Counts per million; FDR, false discovery rate; LR, likelihood ratio.

Supplementary table S9. Differentially expressed surface markers (Cluster of Differentiation molecules). Murine.

Genes shown are censored at FDR P ≤0.05 and log(2) fold change of ±1, and ordered by log fold change. CPM, Counts per million; FDR, false discovery rate; LR, likelihood ratio.

Supplementary table S10. Differentially expressed chemokines. Human.

Genes shown are censored at FDR P ≤0.05 and log(2) fold change of ±1, and ordered by log fold change. Genes highlighted in bold are also significant in the equivalent analysis for murine MAIT cells. CPM, Counts per million; FDR, false discovery rate; LR, likelihood ratio.

Supplementary table S11. Differentially expressed chemokines. Murine.

Genes shown are censored at FDR P ≤0.05 and log(2) fold change of ±1, and ordered by log fold change. Genes highlighted in bold are also significant in the equivalent analysis for human MAIT cells. CPM, Counts per million; FDR, false discovery rate; LR, likelihood ratio.

Supplementary table S12. Differentially expressed chemokine receptors. Human.

Genes shown are censored at FDR P ≤0.05 and log(2) fold change of ±1, and ordered by log fold change. Genes highlighted in bold are also significant in the equivalent analysis for murine MAIT cells. CPM, Counts per million; FDR, false discovery rate; LR, likelihood ratio.

Supplementary table S13. Differentially expressed chemokine receptors. Murine.

Genes shown are censored at FDR P ≤0.05 and log(2) fold change of ±1, and ordered by log fold change. Genes highlighted in bold are also significant in the equivalent analysis for human MAIT cells. CPM, Counts per million; FDR, false discovery rate; LR, likelihood ratio.

Supplementary table S14. Tissue repair gene signature.

Murine tissue repair signature gene set from Linehan *et al*^21^ used in both murine and human GSEA analyses.

Supplementary table S15. Cytometry antibodies.

## References

1. Kjer-Nielsen, L., et al. MR1 presents microbial vitamin B metabolites to MAIT cells. Nature 491, 717–723 (2012).

2. Corbett, A.J., et al. T-cell activation by transitory neo-antigens derived from distinct microbial pathways. Nature 509, 361–365 (2014).

3. Eckle, S.B., et al. Recognition of Vitamin B Precursors and Byproducts by Mucosal Associated Invariant T Cells. J Biol Chem 290, 30204–30211 (2015).

4. Tilloy, F., et al. An invariant T cell receptor alpha chain defines a novel TAP-independent major histocompatibility complex class Ib-restricted alpha/beta T cell subpopulation in mammals. J Exp Med 189, 1907–1921 (1999).

5. Porcelli, S., Yockey, C.E., Brenner, M.B. & Balk, S.P. Analysis of T cell antigen receptor (TCR) expression by human peripheral blood CD4-8-alpha/beta T cells demonstrates preferential use of several V beta genes and an invariant TCR alpha chain. J Exp Med 178, 1–16 (1993).

6. Mak, J.Y., et al. Stabilizing short-lived Schiff base derivatives of 5-aminouracils that activate mucosal-associated invariant T cells. Nature communications 8, 14599 (2017).

7. Le Bourhis, L., et al. Antimicrobial activity of mucosal-associated invariant T cells. Nat Immunol 11, 701–708 (2010).

8. Dusseaux, M., et al. Human MAIT cells are xenobiotic-resistant, tissue-targeted, CD161hi IL-17-secreting T cells. Blood 117, 1250–1259 (2011).

9. Chen, Z., et al. Mucosal-associated invariant T-cell activation and accumulation after in vivo infection depends on microbial riboflavin synthesis and costimulatory signals. Mucosal immunology 10, 58–68 (2017).

10. Wang, H., et al. MAIT cells protect against pulmonary Legionella longbeachae infection. Nature communications 9, 3350 (2018).

11. Meierovics, A., Yankelevich, W.J. & Cowley, S.C. MAIT cells are critical for optimal mucosal immune responses during in vivo pulmonary bacterial infection. Proc Natl Acad Sci U S A 110, E3119–3128 (2013).

12. Meierovics, A.I. & Cowley, S.C. MAIT cells promote inflammatory monocyte differentiation into dendritic cells during pulmonary intracellular infection. J Exp Med 213, 2793–2809 (2016).

13. Reantragoon, R., et al. Antigen-loaded MR1 tetramers define T cell receptor heterogeneity in mucosal-associated invariant T cells. J Exp Med 210, 2305–2320 (2013).

14. Lepore, M., et al. Parallel T-cell cloning and deep sequencing of human MAIT cells reveal stable oligoclonal TCRbeta repertoire. Nature communications 5, 3866 (2014).

15. Jabri, B. & Abadie, V. IL-15 functions as a danger signal to regulate tissue-resident T cells and tissue destruction. Nat Rev Immunol 15, 771–783 (2015).

16. Fabregat, A., et al. The Reactome Pathway Knowledgebase. Nucleic acids research 46, D649–D655 (2018).

17. van Wilgenburg, B., et al. MAIT cells contribute to protection against lethal influenza infection in vivo. Nature communications 9(2018).

18. Heng, T.S., Painter, M.W. & Immunological Genome Project, C. The Immunological Genome Project: networks of gene expression in immune cells. Nat Immunol 9, 1091–1094 (2008).

19. Kanehisa, M. & Goto, S. KEGG: Kyoto encyclopedia of genes and genomes. Nucleic acids research 28, 27–30 (2000).

20. Treiner, E., et al. Selection of evolutionarily conserved mucosal-associated invariant T cells by MR1. Nature 422, 164–169 (2003).

21. Linehan, J.L., et al. Non-classical Immunity Controls Microbiota Impact on Skin Immunity and Tissue Repair. Cell 172, 784–796 e718 (2018).

22. Mootha, V.K., et al. PGC-1alpha-responsive genes involved in oxidative phosphorylation are coordinately downregulated in human diabetes. Nat Genet 34, 267–273 (2003).

23. Kjer-Nielsen, L., et al. MR1 presents microbial vitamin B metabolites to MAIT cells. Nature 491, 717–723 (2012).

24. Godfrey, D.I., Uldrich, A.P., McCluskey, J., Rossjohn, J. & Moody, D.B. The burgeoning family of unconventional T cells. Nat Immunol 16, 1114–1123 (2015).

25. Le Bourhis, L., et al. MAIT Cells Detect and Efficiently Lyse Bacterially-Infected Epithelial Cells. PLoS Pathog 9, e1003681 (2013).

26. Holzapfel, K.L., Tyznik, A.J., Kronenberg, M. & Hogquist, K.A. Antigen-dependent versus-independent activation of invariant NKT cells during infection. J Immunol 192, 5490–5498 (2014).

27. van Wilgenburg, B., et al. MAIT cells are activated during human viral infections. Nature communications 7, 11653 (2016).

28. Loh, L., et al. Human mucosal-associated invariant T cells contribute to antiviral influenza immunity via IL-18-dependent activation. Proc Natl Acad Sci U S A 113, 10133–10138 (2016).

29. Ussher, J.E., et al. TLR signalling in human antigen-presenting cells regulates MR1-dependent activation of MAIT cells. Eur J Immunol 46, 1600–1614 (2016).

30. Rahimpour, A., et al. Identification of phenotypically and functionally heterogeneous mouse mucosal-associated invariant T cells using MR1 tetramers. J Exp Med 212, 1095–1108 (2015).

31. Greene, J.M., et al. MR1-restricted mucosal-associated invariant T (MAIT) cells respond to mycobacterial vaccination and infection in nonhuman primates. Mucosal immunology 10, 802–813 (2017).

32. Klenerman, P. & Ogg, G. Killer T cells show their kinder side. Nature 555, 594–595 (2018).

33. Leng, T., et al. TCR and inflammatory signals tune human MAIT cells to exert specific tissue repair and effector functions. bioRxiv (2018).

34. Rouxel, O., et al. Cytotoxic and regulatory roles of mucosal-associated invariant T cells in type 1 diabetes. Nat Immunol 18, 1321–1331 (2017).

35. Lee, J.S., et al. Interleukin-23-Independent IL-17 Production Regulates Intestinal Epithelial Permeability. Immunity 43, 727–738 (2015).

36. Dudakov, J.A., Hanash, A.M. & van den Brink, M.R. Interleukin-22: immunobiology and pathology. Annu Rev Immunol 33, 747–785 (2015).

37. Sugimoto, K., et al. IL-22 ameliorates intestinal inflammation in a mouse model of ulcerative colitis. J Clin Invest 118, 534–544 (2008).

38. Brown, R.L., Sequeira, R.P. & Clarke, T.B. The microbiota protects against respiratory infection via GM-CSF signaling. Nature communications 8, 1512 (2017).

39. Quinton, L.J., et al. Leukemia inhibitory factor signaling is required for lung protection during pneumonia. J Immunol 188, 6300–6308 (2012).

40. Kurioka, A., et al. Cd161 Defines A Functionally Distinct Subset of Pro-Inflammatory Natural Killer Cells. Frontiers in immunology 9, 486 (2018).

41. Cohen, N.R., et al. Shared and distinct transcriptional programs underlie the hybrid nature of iNKT cells. Nat Immunol 14, 90–99 (2013).

42. Lee, Y.J., et al. Lineage-Specific Effector Signatures of Invariant NKT Cells Are Shared amongst gammadelta T, Innate Lymphoid, and Th Cells. J Immunol 197, 1460–1470 (2016).

43. Eckle, S.B., et al. A molecular basis underpinning the T cell receptor heterogeneity of mucosal-associated invariant T cells. J Exp Med 211, 1585–1600 (2014).

44. Chien, Y.H., Meyer, C. & Bonneville, M. gammadelta T cells: first line of defense and beyond. Annu Rev Immunol 32, 121–155 (2014).

45. Vantourout, P. & Hayday, A. Six-of-the-best: unique contributions of gammadelta T cells to immunology. Nat Rev Immunol 13, 88–100 (2013).

46. Havran, W.L. & Jameson, J.M. Epidermal T cells and wound healing. J Immunol 184, 5423–5428 (2010).

47. Clouthier, D.L. & Watts, T.H. Cell-specific and context-dependent effects of GITR in cancer, autoimmunity, and infection. Cytokine & growth factor reviews 25, 91–106 (2014).

48. Malamut, G., et al. IL-15 triggers an antiapoptotic pathway in human intraepithelial lymphocytes that is a potential new target in celiac disease-associated inflammation and lymphomagenesis. J Clin Invest 120, 2131–2143 (2010).

49. Dobin, A., et al. STAR: ultrafast universal RNA-seq aligner. Bioinformatics 29, 15–21 (2013).

50. Liao, Y., Smyth, G.K. & Shi, W. The Subread aligner: fast, accurate and scalable read mapping by seed-and-vote. Nucleic acids research 41, e108 (2013).

51. Team, R.C. R: A language and environment for statistical computing.. (R Foundation for Statistical Computing, Vienna, Austria, 2014).

52. Robinson, M.D., McCarthy, D.J. & Smyth, G.K. edgeR: a Bioconductor package for differential expression analysis of digital gene expression data. Bioinformatics 26, 139–140 (2010).

53. Yu, G. & He, Q.Y. ReactomePA: an R/Bioconductor package for reactome pathway analysis and visualization. Mol Biosyst 12, 477–479 (2016).

54. G, S., et al. limma: Linear Models for Microarray Data. (2016).

55. Subramanian, A., et al. Gene set enrichment analysis: a knowledge-based approach for interpreting genome-wide expression profiles. Proc Natl Acad Sci U S A 102, 15545–15550 (2005).

56. Maglott, D., Ostell, J., Pruitt, K.D. & Tatusova, T. Entrez Gene: gene-centered information at NCBI. Nucleic acids research 39, D52–57 (2011).

57. Leek, J.T., Johnson, W.E., Parker, H.S., Jaffe, A.E. & Storey, J.D. The sva package for removing batch effects and other unwanted variation in high-throughput experiments. Bioinformatics 28, 882–883 (2012).

58. Gentleman, R.C., et al. Bioconductor: open software development for computational biology and bioinformatics. Genome Biol 5, R80 (2004).

59. Patel, O., et al. Recognition of vitamin B metabolites by mucosal-associated invariant T cells. Nature communications 4, 2142 (2013).

